# Structure-informed direct coupling analysis improves protein mutational landscape predictions

**DOI:** 10.64898/2026.03.27.714804

**Authors:** Matsvei Tsishyn, Hugo Talibart, Marianne Rooman, Fabrizio Pucci

## Abstract

Direct Coupling Analysis has been instrumental over the past decade in leveraging evolutionary information and advancing our understanding of biomolecular structure and function. Here, we introduce sparse extensions of this method that explicitly incorporate structural information. StructureDCA focuses on physically relevant interactions by selectively retaining couplings between residues in spatial contact, and StructureDCA[RSA] additionally incorporates per-residue relative solvent accessibility. These models outperform state-of-the-art approaches in describing mutational landscapes, as they more effectively integrate structural context. Moreover, their sparse formulation enables orders-of-magnitude improvements in computational efficiency while preserving interpretability, providing a powerful framework for gaining mechanistic insights into mutation effects and advancing protein design. The StructureDCA models are available as a user-friendly Python package via the PyPI repository. The source code is freely accessible at https://github.com/3BioCompBio/StructureDCA, which also includes a Colab Notebook interface.

## Introduction

Characterizing the impact of amino acid substitutions in proteins is crucial in many areas of science, biotechnology, and medicine. For instance, it is key in the analysis of genetic variants, which drive evolution but can also cause alterations of protein structure and/or function that lead to pathogenic conditions. Consequently, substantial efforts over the past decades have been devoted to developing computational tools capable of identifying potentially pathogenic variants in human genomes [1]. These tools are fundamental for interpreting genetic variants and complementing clinical evidence [2]. Understanding the effects of amino acid substitutions is also crucial in protein design, where target proteins are rationally modified through selected mutations to improve protein biophysical properties such as stability, solubility, or heat resistance [3, 4].

An important class of computational methods for predicting the effects of amino acid substitutions leverages evolutionary information [5, 6] by exploiting the massive amount of protein sequence data available in public repositories. The underlying idea is to track how evolutionary pressure acts on target proteins by analyzing the multiple sequence alignment (MSA) of their homologs: positions that remain conserved across evolution typically indicate regions with essential structural or functional roles.

MSAs provide information beyond that captured by simple conservation scores. Early studies [7, 8, 9] demonstrated that variation between residue pairs in MSAs provides insight into their spatial proximity. This information can be exploited to predict protein structure and highlights the importance of considering the non-additive effects of mutations, also known as epistasis [10]. Direct coupling analysis (DCA) [11, 12, 13, 14, 15] advances these approaches by representing an MSA as a global statistical model with both positional and pairwise parameters [16]. DCA and related coevolution-based methods have contributed substantially to many biological applications, particularly to residue–residue contact prediction [12, 17, 18, 19], which has in turn inspired the success of protein structure prediction methods such as AlphaFold [20, 21].

Given the importance of epistatic effects in protein sequence evolution [22, 23, 24, 10], DCA methods have also been successfully applied to mutational landscape prediction [25, 26, 27, 28, 29]. However, in our previous work [30], we showed that these methods perform only marginally better than independent-site models on standard mutational benchmarks [30, 31]. We provided evidence that the limited performance gains of coevolutionary models may be related to a higher sensitivity to noise (compared to single-site approaches), owing to their greater complexity and their large number of parameters to be inferred, which often exceeds by far the number of sequences in the MSA.

To address this limitation, coevolutionary models of reduced complexity have been proposed [32, 33]. In particular, a sparse DCA model [34] was developed by iteratively filtering couplings to maximize predictive power on mutational datasets. Although limited by the need for initial mutational data, this approach showed that DCA models with fewer coupling coefficients can outperform their fully connected counterparts.

In this paper, we present a structure-informed approach to DCA. Building on the observation that residues in contact coevolve [35], we reversed the conventional flow of information: whereas DCA has traditionally been used to predict residue–residue contacts [11, 12, 13, 14, 17, 18, 19], we instead leverage known contacts to refine the DCA model (Fig. 1). This significantly reduces model complexity, thereby addressing key shortcomings of standard DCA methods, while enabling substantially faster inference and improved performance. As the field increasingly shifts toward large black-box artificial intelligence approaches, such as protein language models (pLMs) [36], we show that evolution- and physicsbased approaches continue to play a prominent role, achieving comparable or slightly superior performance to state-of-the-art pLMs while preserving simplicity and interpretability.

**Figure 1:**
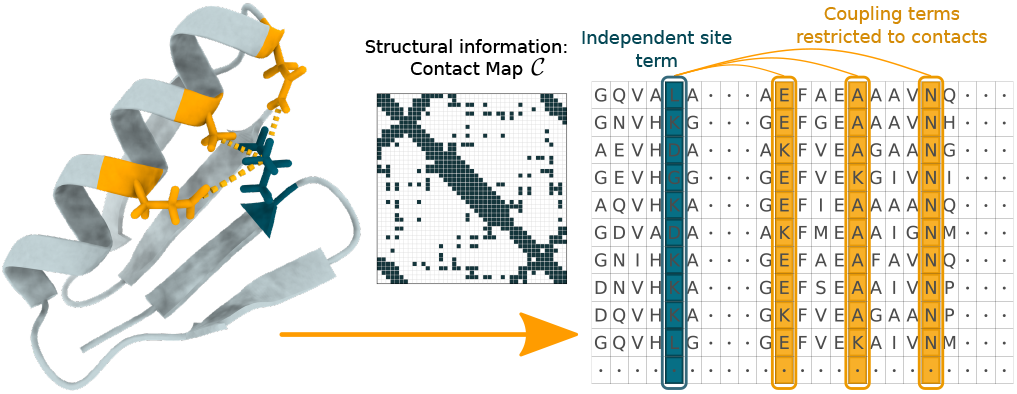
Graphical representation of the StructureDCA model.

## Results

### The StructureDCA model

Standard DCA methods construct a statistical representation of a homologous protein family inferred from its MSA. Let *s* = (*s*_1_, *s*_2_, …, *s*_*L*_) be a protein sequence in the MSA of length *L* and depth *N*, where *s*_*i*_ denotes the amino acid type or a gap at position *i*. The sequence *s* is assumed to have been sampled through evolution with a probability *P* (*s*), described by the Boltzmann distribution, which can be expressed as:

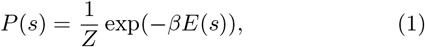

where *β* is the inverse temperature, *Z* denotes the partition function, and *E*(*s*) is the evolutionary energy of *s*, defined as:

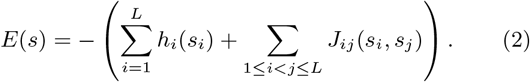

Here, *h*_*i*_(*s*_*i*_) corresponds to single-site fields describing amino acid preferences at individual positions, and *J*_*ij*_ (*s*_*i*_, *s*_*j*_) represents coupling parameters capturing correlations between amino acid types at pairs of positions.

The model parameters *h* and *J* are inferred by maximizing the likelihood of the sequences in the MSA under the probability distribution defined in Eq. 1. Unfortunately, this inference is computationally infeasible. To overcome this issue, several approximation methods have been developed, including mean-field DCA [12], Boltzmann machine DCA [37, 38], and pseudolikelihood maximization DCA [14, 15, 27, 28], which is the method adopted in this study.

To improve the performance of DCA in predicting mutational landscapes, we introduce a new model, named StructureDCA, which generalizes standard DCA by restricting the set of coupling coefficients *J*_*ij*_ . StructureDCA can be viewed as DCA defined on a sparse graphical model with connec-tion graph *C*. The sparsity of this graph is defined based on residue–residue contacts in the three-dimensional (3D) struc-ture of the target protein, which must be provided as input (Fig. 1). This redefines the DCA statistical model so that the energy of a sequence is reformulated as:

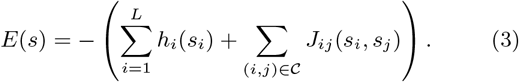

One possible strategy to construct such a sparse DCA model is to first infer the standard fully-connected DCA model and then set to zero all *J*_*ij*_ coefficients corresponding to position pairs not included in the connection graph. However, the coefficients inferred under the fully connected model do not coincide with the pseudolikelihood optimum of the sparse model. For this reason, StructureDCA performs the parameter optimization directly on the restricted parameter space.

Additionally, as shown in our previous studies [30, 31], incorporating the relative solvent accessibility (RSA) of the mutated residue can substantially improve predictions of mutation effects, especially for protein stability. We extend this concept to StructureDCA by introducing, after model inference, positional and pairwise weights on the *h* and *J* contributions. This defines our second model, StructureDCA[RSA]:

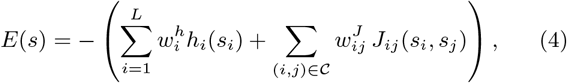

where weights 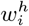 and 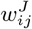 are computed from residues RSA values:

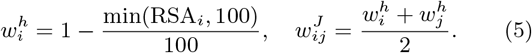

These factors essentially increase the importance of core residues, which are more critical for protein stability than surface residues.

StructureDCA and StructureDCA[RSA] models can be used to predict the effects of amino acid substitutions on biophysical properties by evaluating,

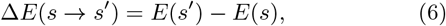

where the difference in statistical energy between the wildtype sequence *s* and the mutant sequence *s*^*′*^ is computed using Eq. 3 for StructureDCA or Eq. 4 for StructureDCA[RSA].

The theoretical framework and implementation of our two models are explained in detail in Supplementary Sections 1 and 2. In Fig. S2, we show an example of computational deep mutational scanning output from StructureDCA and StructureDCA[RSA].

### Improving accuracy and computational efficiency

We started by investigating the relevance of sparsity in DCA models and quantifying its impact on performance in predicting the effects of mutations on protein stability. For this analysis, we explored the full spectrum of sparse DCA models by varying the distance threshold *d*_0_ (ranging from zero to infinity) used to prune coupling coefficients *J*_*ij*_ . This yields a comparison between:

- **Independent-site model** (*d*_0_ = 0): all couplings are removed;
- **StructureDCA** (0 < *d*_0_ < ∞): the retained couplings are determined by the residue–residue contacts in the 3D structure, as defined by the distance threshold *d*_0_;
- **Full DCA** (*d*_0_ = ∞): all couplings are retained, resulting in a standard fully connected DCA model.

We evaluated these models on the MegaScale dataset collection (see Methods), which provides large-scale stability measurements across small protein domains. To assess prediction performance, we computed the average Spearman correlation, *ρ*, between predicted and experimental changes in stability upon mutation across all protein datasets (see Methods and Supplementary Section 3).

As shown in Fig. 2a, the independent-site model and the fully connected DCA model yield similar correlations (*ρ* ≈ 0.48). In contrast, intermediate sparse StructureDCA substantially outperform these baselines, reaching a peak around *d*_0_ = 5 Å where the correlation exceeds *ρ* = 0.54. Performance rises sharply when the first short-range contacts are incorporated into the model (*d*_0_ 4 ∼Å), remains relatively stable up to *d*_0_ = 8 Å, and then gradually declines, likely because additional couplings introduce more noise than useful information. In addition, as shown in Fig. 2a, incorporating RSA into the models consistently improves performance across all distance cutoffs *d*_0_. This shows that structure-guided sparsity and RSA reweighting combine constructively within the model, yielding an overall improvement in correlation from 0.48 up to 0.60. Impressively, the improved performance of StructureDCA over fully connected DCA occurs consistently across almost all individual datasets in the MegaScale collection (Fig. 2b and Figs. S3 and S4).

**Figure 2:**
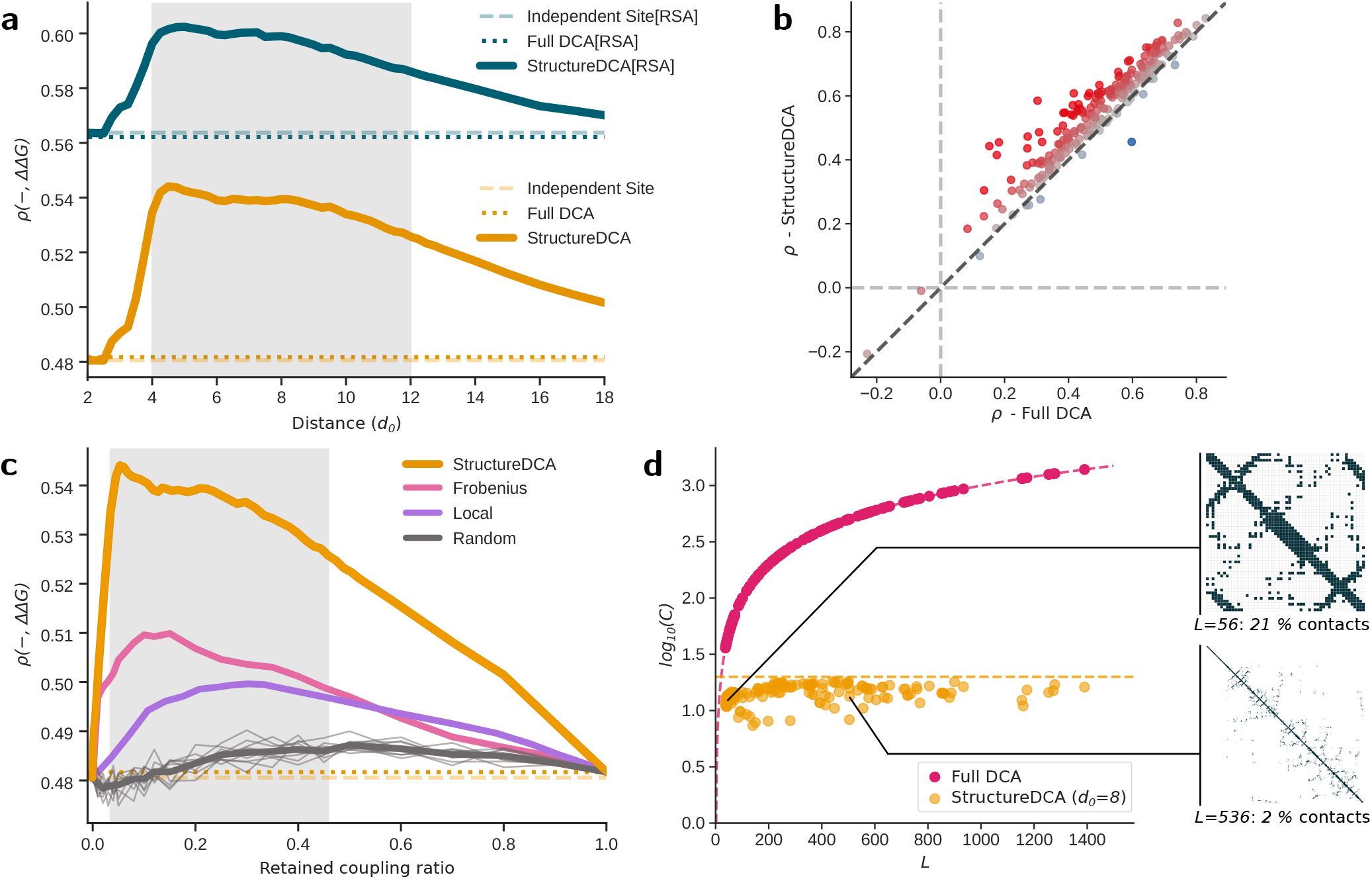
Impact of sparsity on DCA model performance. (a) Average Spearman correlation, *ρ*, on MegaScale for StructureDCA (in orange) and StructureDCA[RSA] (in blue) as a function of the distance cutoff *d*_0_ (in Å). The two extreme baselines, the independentsite model (*d*_0_ = 0Å) and the fully connected DCA model (*d →* ∞) are shown in dashed and dotted lines, respectively. (b) Comparison of Spearman correlations *ρ* for each individual dataset in MegaScale between standard fully connected DCA (*x*-axis) and StructureDCA with *d*_0_ = 8Å (*y*-axis). (c) Average Spearman correlation, *ρ*, on MegaScale for sparse DCA models when the sparsity criterion is the distance cutoff (StructureDCA, orange), the Frobenius norm (pink), the distance along the sequence (purple), and 10 independent random coupling sub-samplings (thin grey lines, with their average in bold line). To enable comparison, all correlations are shown in terms of the retained coupling ratio (on the *x*-axis). (d) Average number of couplings per position, *C* (in log_10_ scale), included in the standard fully connected DCA model (magenta) and in StructureDCA with a distance cutoff *d*_0_ = 8Å (orange), as a function of the MSA length *L*. Target proteins correspond to the ProteinGym dataset, except for two very long proteins that were excluded for clarity but follow the same trend. To illustrate coupling density in StructureDCA, two example contact maps are shown. The first example is a small protein of length *L* = 56, for which 21% of all couplings are retained, and the second is an intermediate-length protein with *L* = 536, for which only 2% of couplings are retained.

To further explore model sparsity, we tested three alternative criteria for constructing sparse DCA models:

- **Frobenius**: keeps only couplings with the strongest coevolutionary signal, as measured by the full DCA Frobenius norm [14];
- **Local**: keeps only couplings between residue pairs that are close in the sequence;
- **Random**: keeps only a set of randomly selected couplings.

Both the Frobenius and local criteria noticeably improve performance relative to the full DCA, with peaks at retained coupling ratios between 0.1 and 0.3 (Fig. 2c). Even random removal of couplings leads to slight improvements. However, these performances remain considerably lower than those achieved using the distance-based criterion.

In summary, only a relatively small number of couplings is sufficient to describe the protein mutational landscape, with couplings between residues in physical contact being the most informative. This approach improves generalizability, enhances robustness to noise, and helps mitigate the undersampling problem [11, 14, 39, 30]. Moreover, the drastic reduction in the number of model parameters, which now grow linearly with protein length instead of quadratically as in fully connected DCA (Fig. 2d), substantially reduces computational time and memory requirements. As shown in Supplementary Section 4, StructureDCA is several orders of magnitude faster than fully connected DCA.

### Comparison with state-of-the-art methods

Here, we show that StructureDCA achieves state-of-the-art performance in predicting the effects of mutations across a wide range of deep mutational scanning (DMS) experiments from three benchmark collections: ProteinGym, MegaScale, and HumanDomains (described in Methods).

We first evaluated StructureDCA on the widely used ProteinGym benchmark (Fig. 3a). Both StructureDCA and StructureDCA[RSA] rank among the top-performing models and are outperformed only by a small number of highly complex pLMs that incorporate 3D structural and/or evolutionary information and comprise hundreds of millions of parameters. In contrast, StructureDCA remains lightweight and interpretable, as its predictions rely on explicit residue-level interactions rather than implicit representations learned by deep neural networks.

**Figure 3:**
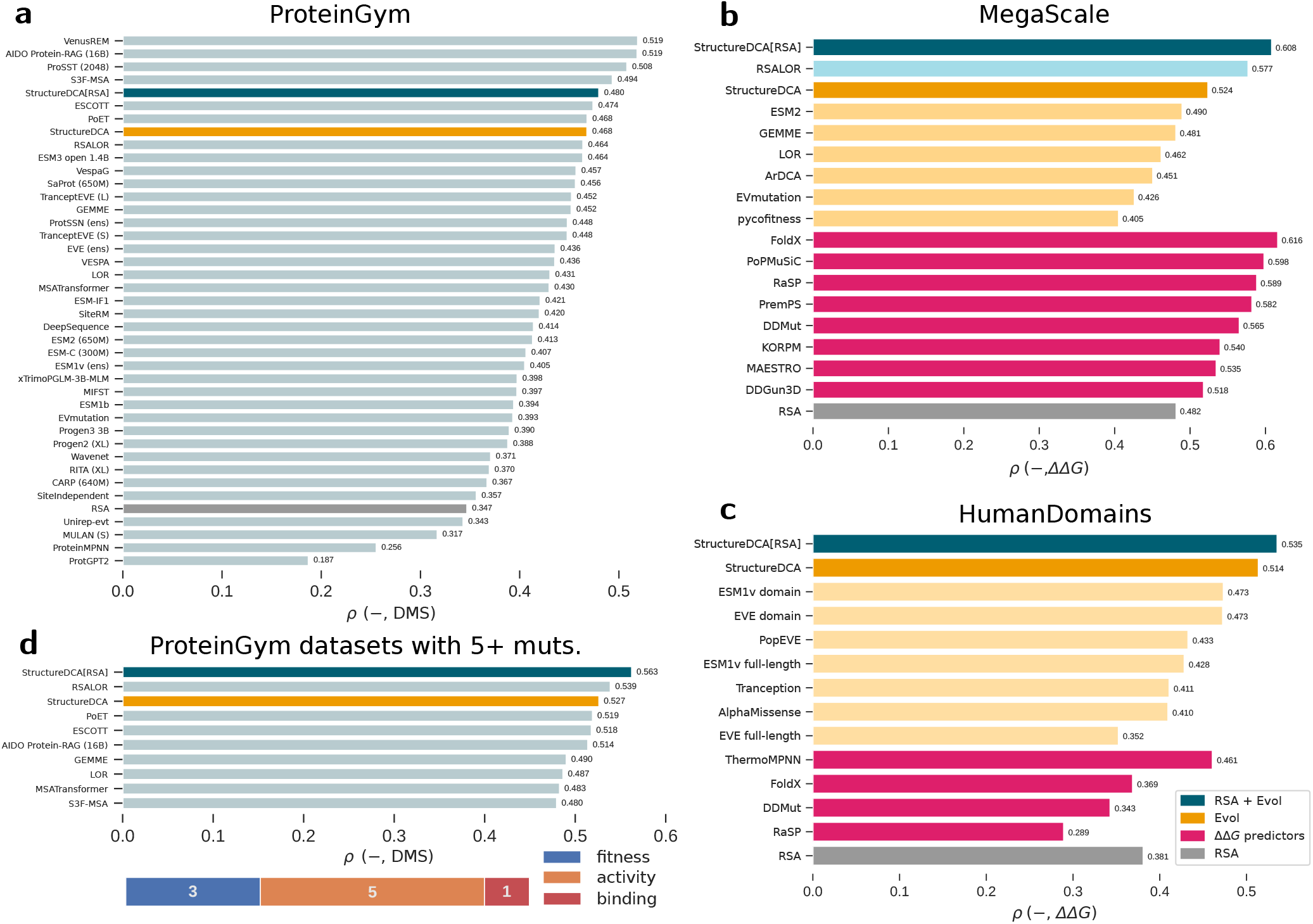
Benchmark of StructureDCA, StructureDCA[RSA], and state-of-the-art methods for predicting mutational effects. Performance is evaluated as the average Spearman correlation, *ρ*, between predictions and experimental measurements. (a) Performance on the ProteinGym dataset collection for methods included in the original benchmark [36], retaining only the best-performing model from each publication. (b) Performance on the MegaScale dataset collection for all methods benchmarked in [30], including classical structure-based ΔΔ*G* predictors (magenta), evolutionary methods (orange), evolutionary methods augmented with RSA (blue), and RSA alone (grey). (c) Performance on the HumanDomains dataset collection for methods benchmarked in [42], using the same color scheme as in (b). (d) Average Spearman correlation across the 9 ProteinGym datasets containing multiple mutations with 5 or more sites mutated simultaneously. Only the bestperforming models are shown. The number of datasets for each DMS type is indicated below. In all panels, StructureDCA, StructureDCA[RSA], and RSA are highlighted in bright orange, blue, and grey, respectively. We also included in the benchmark our independent-site models LOR and RSALOR [31].

Incorporating RSA improves the performance of StructureDCA on ProteinGym, although, as previously reported [31, 40], its relevance depends on the target property measured in the DMS experiment (Fig. S5). Specifically, RSA substantially improves performance on stability datasets, yields modest gains for expression and activity datasets, and slightly reduces performance on fitness datasets. This observation is biologically meaningful, as protein stability primarily depends on core residue–residue interactions, whereas binding and catalytic functions involve partially solvent-exposed residues.

We evaluated our StructureDCA models for predicting protein stability changes upon mutations (ΔΔ*G*) by comparing them with several structure-based ΔΔ*G* predictors (typically trained on literature-based ΔΔ*G* datasets) as well as with unsupervised evolutionary methods, both benchmarked in [30] on the MegaScale dataset collection [41]. StructureDCA outperforms all tested evolutionary models, and StructureDCA[RSA] further extends this lead by outperforming or matching all supervised ΔΔ*G* predictors (Fig. 3b and Fig. S6).

We further evaluated our models on HumanDomains [42], a DMS dataset collection of ΔΔ*G*-related measurements in small human protein domains. Again, StructureDCA and StructureDCA[RSA] outperform all tested models, this time with a notable advance (Fig. 3c and Fig. S6).

Notably, while ΔΔ*G* predictors tend to be better than unsupervised evolutionary methods on MegaScale, the opposite trend is observed on HumanDomains, which is surprising. This suggests that the proxy for stability measurement in the HumanDomains DMS assay may be influenced by fitness-related properties. This interpretation is further supported by the observation that the performance gain provided from incorporating RSA in StructureDCA is smaller on HumanDomains than on MegaScale.

### Capturing epistasis

A first way to evaluate a model’s ability to capture epistatic phenomena is to assess its predictions for higher-order, multisite mutations. A recent study [43] shows that current state-of-the-art models, including pLMs, struggle to predict the non-additive effects of multiple mutations. In this regard, StructureDCA[RSA] markedly enhances performance: it is the top-performing method across the nine ProteinGym datasets containing mutations of order five or higher (Fig. 3d), with similar trends observed for mutations of order 2–4 (Fig. S7).

To further investigate StructureDCA’s ability to capture epistatic effects, we focused on the analysis performed in [44], where experimental DMS was conducted on NDM1 [44] and VIM2 [45], two homologous B1-metallo-*β*-lactamases (MBLs) sharing only ∼ 30% sequence identity. Comparison of the results for these two proteins highlights the effect of the background sequence on their mutational landscapes, reflecting underlying epistatic interactions.

We constructed a single StructureDCA([RSA]) model using the MSA of the MBL family provided in the original publication, excluding the NDM1 and VIM2 sequences, and incorporating contact maps and RSA values extracted from both NDM1 and VIM2 experimental structures (Fig. 4a). By using a single StructureDCA([RSA]) model to predict the mutational landscapes of both proteins, we ensured that the observed differences arise solely from the background sequence rather than from differences in the input data.

**Figure 4:**
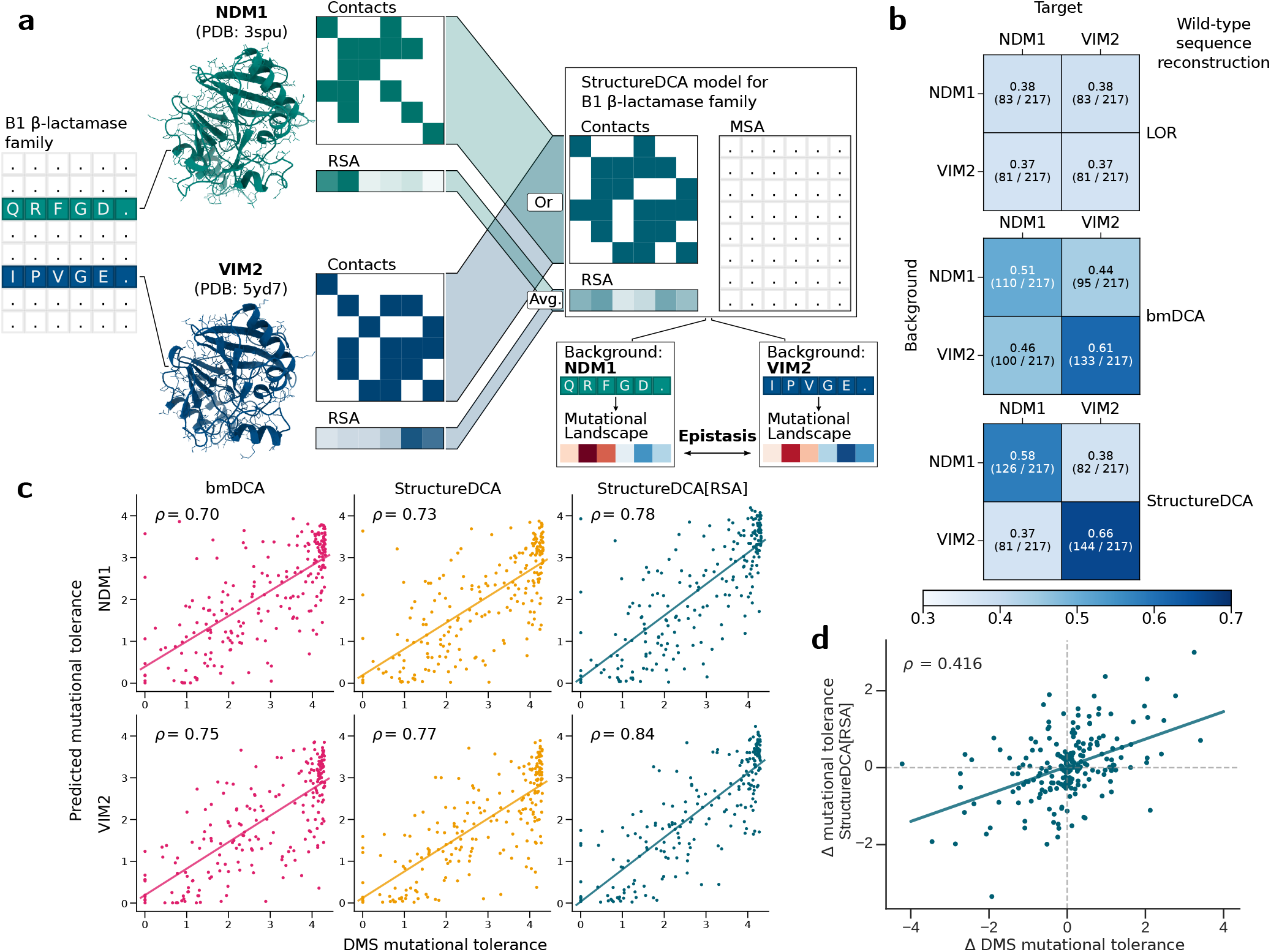
Epistasis between NDM1 and VIM2 B1 metallo-*β*-lactamases. (a) Workflow for quantifying epistasis between NDM1 and VIM2 using StructureDCA and StructureDCA[RSA]. A single StructureDCA([RSA]) model was built using the experimental structures of both proteins (PDB codes 3spu and 5yd7 [46]). Residue contacts were defined when present in at least one structure. RSA values were averaged over the two protein structures (Fig. S8). The mutational landscapes were evaluated relative to each target sequence, NDM1 and VIM2, so that observed differences between the landscapes arise solely from background epistatic effects. (b) The NDM1 and VIM2 target sequences were reconstructed by masking each position individually and predicting the most favorable amino acid based on the relative energy changes Δ*E* as predicted by LOR, bmDCA and StructureDCA. Predictions were performed using either the native sequence background or the alternative background (among NDM1 or VIM2). Each cell reports the reconstruction accuracy rate and the count of correctly recovered positions. (c) Comparison of mutational tolerance between experimental DMS values and predicted values using bmDCA (magenta), StructureDCA (orange) and StructureDCA[RSA] (blue). (d) Comparison of the mutational tolerance difference (NDM1− VIM2) between experimental DMS measurements and predictions using StructureDCA[RSA]. In (c) and (d), the lines represent linear regressions, and the corresponding Spearman correlations *ρ* are shown.

We first compared the experimental DMS measurements with the Δ*E* values obtained from Boltzmann machine-based DCA (bmDCA) [37, 38], as performed in the original study, as well as with predictions from StructureDCA and StructureDCA[RSA] (details in Fig. S8). All three DCA methods reproduce the experimental data well, with StructureDCA slightly outperforming bmDCA. Performance is further improved by StructureDCA[RSA], reaching Spearman correlations of *ρ* = 0.71 and *ρ* = 0.77 for NDM1 and VIM2, respectively (Fig. S8).

To assess the importance of epistatic effects, we performed a sequence reconstruction experiment. For each sequence, every amino acid was masked in turn and reconstructed as the lowest-energy amino acid using either the corresponding wild-type background sequence (e.g., reconstructing NDM1 using the NDM1 background) or the alternative background sequence (e.g., reconstructing the NDM1 sequence using the VIM2 background) (Fig. 4b). As a baseline, our independentsite model LOR, which ignores epistatic effects, reconstructs the same sequence regardless of the background and achieves a relatively low reconstruction accuracy rate of 0.38. In contrast, both bmDCA and StructureDCA produce substantially more accurate reconstructions when the correct background is used. Notably, StructureDCA exhibits markedly stronger background sensitivity: reconstructions are more accurate with the wild-type background but worse when using the alternative background. StructureDCA[RSA] provides similar reconstruction results to StructureDCA.

In addition, we predicted the mutational tolerance at each position of the two MBL proteins (Supplementary Section 1.9), which quantifies the extent to which a given position can accept mutations while maintaining protein fitness. Notably, all tested DCA approaches accurately identified the experimentally defined sensitive positions (Fig. 4c). Since mutational tolerance is independent of the wild-type amino acid at a given position, these values can be compared between NDM1 and VIM2 even at positions where the sequences differ, enabling a more detailed analysis of epistatic contributions at each site. Experimentally, large differences in tolerance are observed at some positions between the two proteins, reflecting their distinct mutational landscapes. Strikingly, these differences are well captured by the predictions, particularly by StructureDCA[RSA] (Fig. 4d and Fig. S9).

Overall, across this computational experiment, StructureDCA consistently outperformed bmDCA, with further improvements achieved using StructureDCA[RSA]. Moreover, StructureDCA and StructureDCA[RSA] dramatically reduced computation time by approximately three orders of magnitude compared with bmDCA (`adabmDCA 2.0`). This impressive gain in computational efficiency greatly expands the scale of feasible experiments, enabling analyses at the proteome scale.

### Capturing coevolution in PPIs

We next demonstrate that StructureDCA effectively captures (co)evolutionary signals in protein–protein interactions (PPIs) and integrates structural information to generate results consistent with biological interpretation.

We first considered the DMS experiment from [47], which focuses on four sites in the ParD antitoxin of a prokaryote species and performs combinatorial mutagenesis, encompassing single, double, triple, and quadruple substitutions at these positions (Fig. S10). The experimental assay measures organismal fitness, which is strongly related to the ability of ParD to bind its cognate toxin protein, ParE. For a complete analysis of StructureDCA predictions on this dataset, we analyzed subsets of lower-order mutations separately (see explanations in Fig. S10).

On this dataset, when using the MSA provided by ProteinGym, StructureDCA exhibits relatively modest accuracy compared with its performance on other ProteinGym datasets, placing it among the lowest-performing 5%. The Spearman correlations between predicted and experimental fitness values are *ρ* = 0.11 for single and double mutants, *ρ* = 0.19 for mutations up to order 3, and *ρ* = 0.18 for the full dataset. These low scores can be partially explained by the monomeric nature of the input structure provided by ProteinGym, in which the four mutated positions lie within the same same *α*-helix and form only a few local intra-chain contacts (Fig. 5a, and Fig. S11 for additional details). In fact, these four residues are located at the center of the ParD–ParE interface and establish multiple contacts with residues of the ParE toxin. These residues were originally selected for mutagenesis because of their strong coevolutionary signal with the toxin [47]. In line with this observation and previous find-ings [27], the performance of StructureDCA improves substantially (Fig. 5a) when using a concatenated ParD–ParE MSA and deriving the contacts from the experimental structure of the ParD–ParE complex. The Spearman correlations increase to *ρ* = 0.64 for single and double mutations, *ρ* = 0.57 for mutations up to order 3 and *ρ* = 0.39 on the full dataset. Remarkably, removing all intra-chain couplings, thus considering only inter-chain interactions, further improves agreement between the coevolutionary model and experimental data (Fig. 5a and Fig. S12).

**Figure 5:**
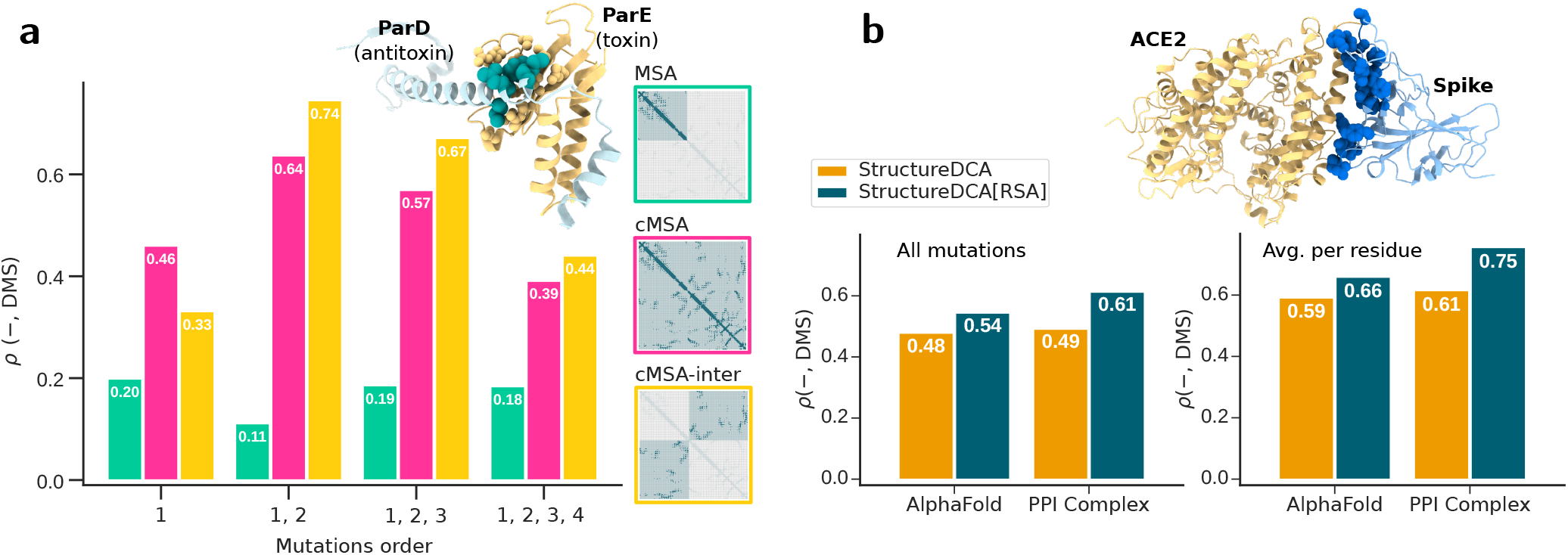
Protein–protein interactions. (a) ParD–ParE (toxin–antitoxin complex). Spearman correlations, *ρ*, between StructureDCA predictions and experimental DMS data [47] using the MSA of the ParD monomer (green, “MSA”), the concatenated ParD-ParE MSA (magenta, “cMSA”), and the concatenated ParD-ParE MSA considering only inter-chain couplings (yellow, “cMSA-inter”). Performance is evaluated on subsets containing increasing mutation orders (i.e., number of mutations), up to the full dataset. StructureDCA predictions are computed using the experimental ParD–ParE structure (PDB code `5ceg` [47]). Additional details on StructureDCA runs are provided in Fig. S11. The four residues mutated in the DMS dataset are highlighted as blue spheres on the 3D structure. (b) SARS-CoV-2 Spike RBD–human ACE2 interaction. Spearman correlations, *ρ*, between StructureDCA and StructureDCA[RSA] predictions and experimental DMS data [48], computed using either the AlphaFold monomeric structures or the experimental dimeric structure (PDB code `6m0j` [49]). Results are provided for all mutations (left) and for per-residue averaged values (right). Residues whose RSA is affected by the interaction are highlighted as blue spheres on the 3D structure.

The second case study concerns the effect of single-site mutations on the binding of the SARS-CoV-2 spike receptor-binding domain (RBD) to the human ACE2 receptor [48]. Because this is an inter-species interaction, coevolutionary signals between the two proteins cannot be directly exploited: while the spike protein is under strong evolutionary pressure to bind ACE2, this is not the case for the human receptor.

Moreover, ProteinGym provides a monomeric 3D structure for this entry, whereas the bound conformation is more biologically relevant. Using the experimental 3D structure of the complex (PDB code `6m0j` [49]) instead of the monomeric model substantially improves predictive performance. The Spearman correlation between prediction and experimental values increases from *ρ* = 0.54 to *ρ* = 0.61 for all mutations, and from *ρ* = 0.66 to *ρ* = 0.75 when averaging per residue (Fig. 5b), making StructureDCA[RSA] the best-performing model for this protein among all methods benchmarked in ProteinGym. A similar trend is observed for another PPI (Fig. S13).

## Discussion

DCA methods have played a prominent role in extracting coevolutionary signals from MSAs of homologous families, enabling advances in biomolecular structure prediction and functional characterization. Recently, however, AI-based approaches such as pLMs have gained increasing interest by outperforming traditional evolutionary models [36]. We showed here that integrating structural information to DCA models significantly enhances the accuracy of protein mutational landscape predictions and achieves performances comparable to, or slightly exceeding, that of state-of-the-art AI models. Additionally, the drastic reduction in the number of parameters makes our model orders of magnitude faster than standard DCA, enabling proteome-scale evolutionary analyses.

Importantly, beyond gains in speed and predictive accuracy, StructureDCA and StructureDCA[RSA] remain substantially more interpretable than black-box machine learning models, thus providing valuable mechanistic insights. Namely, we showed that restricting the model to biophysically relevant couplings between residues yields a more accurate description of protein evolution, whereas retaining all couplings tends to introduce noise. While epistatic contributions are known to play a central role in shaping mutational landscapes, current state-of-the-art methods often fail to accurately capture them [43], whereas StructureDCA([RSA]) provides meaningful insights into epistasis. Finally, we highlight the importance of incorporating the correct structural context, which is often overlooked by AI-based models but effectively captured by StructureDCA([RSA]).

The StructureDCA([RSA]) model is provided as an easy-to-install, user-friendly tool, available both as a PyPI package and via a Colab Notebook. This offers flexible parameter customization and enables broad use within the scientific community, including users without bioinformatics expertise. While this study primarily focuses on predicting mutational landscapes, StructureDCA and StructureDCA[RSA] can also perform a range of additional analyses, such as customizing model couplings to construct sparse DCA models tailored to specific biological contexts.

## Methods

### Mutational datasets

To assess the performance and robustness of StructureDCA and StructureDCA[RSA] in predicting protein mutational landscapes across diverse applications, we considered a large set of protein DMS datasets:

- **ProteinGym** [36] is a standard and widely used benchmark dataset collection designed to test and validate unsupervised models that predict mutational effects. It comprises a diverse set of 217 DMS datasets covering both single-site and multiple mutations across a broad range of protein biophysical properties, including stability, binding affinity, organismal fitness, activity, and expression. To ensure a fair comparison with other methods in the ProteinGym benchmark, we used the dataset collection without any modifications.
- **MegaScale** [41] contains DMS datasets that evaluate the effects of single-site and multiple mutations on protein thermodynamic stability (ΔΔ*G*) for 412 small protein domains. It provides a high quality, systematic and large-scale set of measurements, all performed with the same experimental setup in a “single” cell-free cDNA display proteolysis experiment. We removed *de novo* proteins and proteins with low-depth MSAs (fewer than 100 unique sequences), yielding a final set of 238 protein domains. Note that most of the ProteinGym DMS experiments measuring stability come from the MegaScale dataset.
- **HumanDomains** [42] is a DMS dataset collection similar to MegaScale, but measuring the effects of mutations on protein thermodynamic stability (ΔΔ*G*) using an abundance protein fragment complementation assay. It contains only single-site mutations in 522 small human protein domains. We discarded proteins with a very lowdepth MSA (less than 100 sequences), resulting in a final set of 509 protein domains.

### MSA and 3D structure curation

For the ProteinGym benchmark, we used the MSAs provided in the original publication [36]. For the MegaScale and HumanDomains datasets, we generated the MSAs using EVcouplings [50] (with default parameters), which relies on JackHMMER [51] to build an initial MSA and applies additional post-processing steps. We used UniRef90 [52] as protein sequence database. For the MBL family, we used the MSA provided in the original publication [44]. The concatenated MSA of the ParD–ParE system was obtained using the ColabFold pipeline [53].

MSAs usually contain gaps due to insertions and deletions. Some DCA methods treat gaps as another type of amino acid [11, 12]. Although this option is available and can be useful for analyzing sequences containing deletions, gap characters are excluded by default from the StructureDCA coefficient optimization step, as in EVmutation. [27]. This avoids assigning excessive importance to correlations involving highly gapped positions.

Moreover, sequences in an MSA are unevenly distributed across sequence space, leading to redundancy among closely related sequences. To address this issue, the relative contributions of sequences from the MSA are reweighted [54, 11, 12] by reducing the impact of populated clusters of highly similar sequences (by default, more than 80% sequence identity), as we did in [30, 31].

Finally, MSAs often contain sequences in the “twilight zone” [55], i.e., sequences that are too evolutionarily distant from the target sequence based on sequence identity criteria. Following our previous work [30, 31], such sequences (by default, those with less than 25% sequence identity to the target) are excluded from the MSA.

As structural information for the different computational benchmarks, we used the monomeric protein 3D structures generated by AlphaFold [21], as provided in the original publications. For the application of StructureDCA to ParD–ParE complex, spike protein–ACE2 complex, and the two MBL proteins, we used experimental structures retrieved from the Protein Data Bank [56].

### Evaluation metrics

The primary evaluation metric is Spearman’s rank correlation coefficient, *ρ*, computed between predicted and experimental values, enabling comparison across datasets with different scales. For each collection of DMS datasets, we computed the Spearman correlation separately for each dataset and reported the average value. For ProteinGym and MegaScale, the contributions of individual datasets were reweighted to account for the presence of highly similar target proteins (Supplementary Section 3).

### Additional functionalities of StructureDCA

Beyond their application to mutational analysis in this study, StructureDCA and StructureDCA[RSA] support a wide range of additional tasks. In particular, they enable the evaluation of the energy of a target sequence under a model inferred from a multiple sequence alignment (MSA), as well as access to model parameters, including the fields *h*, pairwise couplings *J*, and their associated Frobenius norms. They integrate a high-performance standard and sparse DCA solver (implemented in C++) with a user-friendly and highly flexible Python interface, exposing a wide range of customiz-able parameters. These parameters cover key steps such as MSA curation, regularization of DCA coefficients, and residue–residue distance determination.

Furthermore, StructureDCA allows the construction of sparse DCA models using user-defined contact maps, rather than relying on the default distance-based criteria, and StructureDCA[RSA] enables the use of custom weights instead of the default RSA-based weights. This flexibility permits the derivation of a wide variety of sparse DCA models tailored to specific applications.

Additional details are provided in Supplementary Section 2 and in the GitHub repository, and a more comprehensive analysis of these aspects will be presented in a forthcoming publication.

## Data and code availability

All datasets, experimental measurements, metadata and StructureDCA predictions are available in our data repository https://zenodo.org/records/18955525.

StructureDCA and StructureDCA[RSA] are free and open source (source code: https://github.com/3BioCompBio/StructureDCA). They can be installed as a Python package (PyPI) with a single command `pip install structuredca.`

They are also available via a Colab Notebook providing a user-friendly graphical interface that automatically retrieves or generates the MSA and the 3D structure of the target protein, infers the StructureDCA and StructureDCA[RSA] models, and performs mutational analyses or sequence energy evaluations. The predicted mutational landscape can also be visualized as a heatmap and mapped onto the 3D structure.

## Acknowledgments

We acknowledge financial support from the F.R.S.-FNRS Fund for Scientific Research. M.T. has benefited from F.R.S.-FNRS FRIA and PDR PhD grants, and H.T. from a F.R.S.-FNRS PDR postdoc grant. Computational resources were provided by the Consortium des Équipements de Calcul Intensif (CÉCI), funded by the Walloon Region and the F.R.S.-FNRS. We also thank Martin Weigt, Matteo Bisardi, Alessandra Carbone, and Elodie Laine for insightful discussions and comments on the topic.

## Notes

### Competing Interest Statement

The authors have declared no competing interest.

