## Supplementary Material for "Structure-informed direct coupling analysis improves protein mutational landscape predictions"

Matsvei Tsishyn[1,2], Hugo Talibart[1,2], Marianne Rooman[1,2], Fabrizio Pucci[1,2]
[1] Computational Biology and Bioinformatics, Université Libre de Bruxelles, 1050 Brussels, Belgium
[2] Interuniversity Institute of Bioinformatics in Brussels, 1050 Brussels, Belgium

March 27, 2026

**3 Evaluation metrics and datasets** **12**

**4 StructureDCA computational time complexity and efficiency** **13**

**5 Results: Supplementary Tables and Figures** **15**

### 1 Theoretical foundations of StructureDCA

#### 1.1 Multiple sequence alignments

The primary input to direct coupling analysis (DCA) models is a multiple sequence alignment (MSA) representing a family of $N$ homologous protein sequences. Sequences are aligned by introducing gaps so that all sequences have the same length, $L$, and functionally homologous residues across different sequences are aligned in the same positions (columns) as much as possible. An MSA is therefore a representation of the evolutionary history of a family of homologous protein sequences that have been subject to similar evolutionary pressures.

Formally, we represent an MSA of length $L$ with $N$ sequences as a 2-dimensional array,

$$\mathbf{S} = (s^n)_{n=1}^{N}, \tag{1}$$

where $s^n$ is the $n^{\text{th}}$ sequence of the MSA. Each sequence $s = (s_1, \ldots, s_L)$ is an array of symbols on an alphabet of length $q = 21$ (20 standard proteinogenic amino acids and 1 gap symbol). The state (amino acid type) in the $n^{\text{th}}$ sequence at position $i$ it noted as $s_i^n$.

#### 1.2 Positional frequencies

We compute the frequency of an amino acid $a \in \{1, \ldots, q\}$ at a position $i \in \{1, \ldots, L\}$ in the MSA as,

$$f_i(a) = \frac{1}{N} \sum_{n=1}^{N} \delta_{a\, s_i^n}, \tag{2}$$

where $\delta$ is the Kronecker delta, which equals one if the two indices are equal and zero otherwise. The single site frequency $f_i$ can be interpreted as an indicator of the sequence conservation at position $i$. For instance, a functionally important, highly conserved position $i$ can have a frequency $f_i(a)$ close to one for a given amino acid $a$ while all other frequencies are close to zero. On the other hand, a variable position (for instance in a loop), could display a more evenly distribution of frequencies.

Similarly, we can also derive coupled frequencies for a pair of amino acids $a$ and $b$ at positions $i$ and $j$ respectively as,

$$f_{ij}(a, b) = \frac{1}{N} \sum_{n=1}^{N} \delta_{a\, s_i^n} \cdot \delta_{b\, s_j^n}. \tag{3}$$

An idealized example often used to illustrate the importance of coupled frequencies $f_{ij}$ is an evolutionary conserved salt bridge between two residues at positions $i$ and $j$, respectively. Here, a positively charged amino acid occupies position $i$ whenever a negatively charged amino acid occupies position $j$, and conversely. In this scenario, single-site frequencies are thus not sufficient to describe the evolutionary pressure at positions $i$ and $j$.

#### 1.3 Weighting and regularization

Inferring a model from an MSA assumes that the sequences it contains are independent samples drawn from a probability distribution describing the homologous protein family. However, phylogenetic relationships between species can result in highly similar sequences that have not yet had sufficient time to diverge. This effect is amplified by the selection bias of sequenced species in biological databases. These sampling biases can introduce correlations that do not accurately describe structural and functional properties of the protein family [1].

A common method to mitigate this issue is to assign a weight factor to each sequence of the MSA in order to downgrade the importance of overly represented clusters of sequences [2]. In this study, the weight $w^n$ of a sequence $s^n$ in an MSA $\mathbf{S}$ is defined as the inverse of the number of sequences in the MSA sharing a sequence identity with $s^n$ greater than or equal to a threshold value $\tau$:

$$w^n = \frac{1}{|\{s \in \mathbf{S} \mid \mathrm{seqid}(s^n, s) \geq \tau\}|}. \tag{4}$$

A typical sequence identity threshold is $\tau = 0.80$. These weights define $N_{\mathrm{eff}}$, the effective number of sequences in the MSA [3] as,

$$N_{\mathrm{eff}} = \sum_{n=1}^{N} w^n. \tag{5}$$

As $N_{\mathrm{eff}}$ more accurately represents the number of independent sequences than $N$, it can be used as a measure of the amount of available sequence information in the MSA [4].

On the other hand, the fact that an amino acid $a$ is never observed at a position $i$ (so that $f_i(a) = 0$) does not necessarily mean that this configuration is impossible. Indeed it can simply be a consequence of the fact that the number of sequences in the MSA is limited or that evolution has not explored this possibility yet. To address this issue, we apply a standard regularization approach by introducing a pseudocount $\theta$ into the empirical frequency estimates [5]:

$$f_i(a) \to (1-\theta) f_i(a) + \frac{\theta}{q}. \tag{6}$$

Incorporating weighting and frequency regularization, we reformulate the definition of single-site and coupled frequencies given in Eq. 2 and 3 as

$$f_i(a) = \left( \frac{1-\theta}{N_{\mathrm{eff}}} \sum_{n=1}^{N} w^n \delta_{a\, s_i^n} \right) + \frac{\theta}{q}, \tag{7}$$

$$f_{ij}(a,b) = \left( \frac{1-\theta}{N_{\mathrm{eff}}} \sum_{n=1}^{N} w^n \delta_{a\, s_i^n} \cdot \delta_{b\, s_j^n} \right) + \frac{\theta}{q^2}. \tag{8}$$

##### 1.4 Potts model and DCA

Borrowed from statistical physics, the generalized Potts model is a mathematically well-defined formalism that can statistically describe a family of homologous protein sequences in an MSA [6]. We assume that the sequences in the MSA constitute a sample from a Boltzmann distribution and that the underlying probability distribution is completely determined by single-site fields and pairwise couplings (ignoring all higher order terms). Although this is a simplification of biological reality, it appears to be a useful way to extract information from sequence data [1].

In contrast to independent statistics, such as mutual information derived from single-site and pairwise MSA frequencies (Eqs 2 and 3), the aim of the generalized Potts model is to define a global statistical model of the MSA. A generalized Potts model that describe an MSA is called DCA model [3, 7, 8, 9, 1]. Concretely, a DCA model of length $L$ is parameterized by field coefficients $h_i(a)$, describing the preference for amino acid $a$ at position $i$ in the MSA, and coupling coefficients $J_{ij}(a,b)$, describing the preference for the joint occurrence of amino acids $a$ and $b$ at positions $i$ and $j$, respectively. These coefficients define the evolutionary statistical energy $E$ of a sequence $s$ of length $L$ within the context of the corresponding MSA as:

$$E(s) = -\left( \sum_{i=1}^{L} h_i(s_i) + \sum_{1 \leq i < j \leq L} J_{ij}(s_i, s_j) \right). \tag{9}$$

This expression is the Hamiltonian of the system and its explicit form is sufficient to completely define the system, as all statistical properties can be derived from it.

The probability of a sequence $s$ of length $L$ being sampled from this MSA is then given, by definition of the Boltzmann distribution, as:

$$P(s) = \frac{1}{Z} \exp\left(\sum_{i=1}^{L} h_i(s_i) + \sum_{1\leq i<j\leq L} J_{ij}(s_i, s_j)\right), \tag{10}$$

where the partition function $Z$ normalizes the sum of probabilities over all possible sequences of length $L$ to 1. Note that the information contained in the pair of positions $i$ and $j$ is identical to that in the pair $j$ and $i$; in other words, position pairs are considered unordered. The couplings $J_{ij}$ are thus only identified for $i < j$ in the model. For convenience, we define $J$ as a $L \times L$ matrix with the following relations:

$$J_{ij}(a, b) = \begin{cases} J_{ji}(b, a) & i > j \\ 0 & i = j. \end{cases} \tag{11}$$

Finally, the difference in evolutionary energy, $\Delta E$, of a mutation at position $i$ in a sequence $s$ from wild-type amino acid $a$ to a mutant amino acid $b$ is as derived from Eqs 9 and 11:

$$\begin{aligned} \Delta E(i, a \to b \mid s) &= E(s_1, \ldots, s_{i-1}, b, s_{i+1}, \ldots, s_L) - E(s_1, \ldots, s_{i-1}, a, s_{i+1}, \ldots, s_L) \\ &= -\Delta h_i(a \to b) - \sum_{j=1}^{L} \Delta J_{ij}(a \to b, s_j) \\ &= h_i(a) - h_i(b) + \sum_{j=1}^{L} (J_{ij}(a, s_j) - J_{ij}(b, s_j)). \end{aligned} \tag{12}$$

This equation easily generalizes to multiple mutations.

#### 1.5 Inferring Potts model coefficients

The inference of the coefficients, sometimes referred to as the inverse Potts problem, is performed empirically using the set of sequences in the MSA. The goal is to find the coefficients $\{h, J\}$ that best describe the observed sequences according to the maximum likelihood principle (MLP). More formally, given an MSA of length $L$ with $N$ sequences $\mathbf{S} = (s^n)_{n=1}^{N}$, we seek the coefficients $\{h, J\}$ that maximize the log-likelihood that the specific sequences from the MSA were sampled:

$$\mathcal{L}(h, J) = -\sum_{n=1}^{N} \log P(s^n). \tag{13}$$

In principle, since $\mathcal{L}$ is differentiable, one could solve the system of equations $\partial h_i(a)\mathcal{L} = 0$ and $\partial_{J_{ij}(a,b)}\mathcal{L} = 0$ to explicitly determine the values of the optimal coefficients. However, this approach is computationally intractable, as it requires evaluating the partition function $Z$, which involves summing over all $q^L$ possible sequences of length $L$. For this reason, empirical inference of generalized Potts model parameters typically relies on approximations and remains a challenging and active area of research [6].

#### 1.6 Pseudo-likelihood approximation

One possible approach to solving the above-described MLP problem (Eq. 13) is to use the pseudo-likelihood maximization approximation (PLM). The idea is to approximate the probability of a sequence $s$ using conditional probabilities, thus removing the need to evaluate the partition function $Z$. This approach is widely used as it provides a good trade-off between computational efficiency and precision. In addition, it appears to be relatively easy to adapt to the case of sparse DCA models (as shown in the next section).

We first introduce the conditional probability $P_i^n(a)$ of observing amino acid (or gap) $a$ in sequence $s^n$ at position $i$, given amino acids (or gaps) at all other positions $s^n_{\setminus i} \coloneqq (s_1^n, \dots, s_{i-1}^n, s_{i+1}^n, \dots, s_L^n)$, which is defined as:

$$P_i^n(a) \coloneqq P(s_i = a \mid s^n_{\setminus i}) = \frac{\exp\left(h_i(a) + \sum_{j=1}^{L} J_{ij}(a, s_j^n)\right)}{\sum_{b=1}^{q} \exp\left(h_i(b) + \sum_{j=1}^{L} J_{ij}(b, s_j^n)\right)}. \tag{14}$$

The full probability of sequence $s^n$ is then approximated as the product, over all positions, of conditional probabilities:

$$P(s^n) \to \prod_{i=1}^{L} P_i^n(s_i^n). \tag{15}$$

Note that this procedure is similar to the masking process in large language models, where each token is masked and then predicted using all other tokens in the sequence. Using this approximation, we formulate the pseudo-likelihood to be maximized as:

$$\begin{aligned} \mathcal{PL}(h, J) &= -\sum_{n=1}^{N} \log \prod_{i=1}^{L} P_i^n(s_i^n) \\ &= -\sum_{n=1}^{N} \sum_{i=1}^{L} \log P_i^n(s_i^n). \end{aligned} \tag{16}$$

Analogous to other pseudo-likelihood DCA implementations, we include $L2$ regularization terms in the previous equation to mitigate PLM overfitting toward represented sequences in a given MSA and to prevent divergence of the coefficients. Specifically, we penalize large $h$ and $J$ parameters using two distinct L2 regularization factors, $\lambda_h$ and $\lambda_J$. In addition, we reweight the contribution of each sequence $s^n$ in the MSA by its weight $w^n$ (Eq. 4). Including $L2$ regularization and sequence reweighting, the PLM equation (Eq. 16) becomes:

$$\mathcal{PL}^{L2,w}(h, J) = -\sum_{n=1}^{N} w^n \left( \sum_{i=1}^{L} \log P_i^n(s_i^n) \right) + \lambda_h \sum_{i=1}^{L} \sum_{a=1}^{q} h_i(a)^2 + \lambda_J \sum_{i,j=1}^{L} \sum_{a,b=1}^{q} J_{ij}(a, b)^2. \tag{17}$$

This adapted PLM equation can be solved numerically using a gradient descent approach.

##### 1.7 Sparse Direct Couplings Analysis

In contrast to standard fully connected DCA, where all pairs $\{i, j\}$ are associated with a coupling matrix $J_{ij}$, the model can be generalized to a sparse formulation by defining only a subset of these couplings. The sparsity of the model is then defined by a binary connection graph $\mathcal{C}$ on the set $\{1, \dots, L\}$, where $L$ is the length of the MSA. Equivalently, $\mathcal{C}$ can be viewed as a symmetric binary connection matrix of shape $L \times L$ with zero diagonal (as self-connections are never allowed for $J$ coefficients). Interestingly, a sparse DCA model can represent the full spectrum from a profile-based model, considering only positional amino acid frequencies (when $\mathcal{C}$ is empty), to the fully connected DCA model (when $\mathcal{C}$ is full).

In this case, the Hamiltonian (Eq. 9) is reformulated as

$$E(s) = -\left( \sum_{i=1}^{L} h_i(s_i) + \sum_{i<j \mid (i,j) \in \mathcal{C}} J_{ij}(s_i, s_j) \right). \tag{18}$$

The probability of a sequence $s$ and all other DCA-derived quantities are adapted similarly. Notably, the pseudo-likelihood maximization (Eq. 17) formalism can be trivially generalized to sparse DCA models by removing couplings $J_{ij}$ not included in the connection graph $\mathcal{C}$. The effect of a single-site mutation can be expressed by adapting Eq. 12,

$$\Delta E(i, a \to b \mid s) = -\Delta h_i(a \to b) - \sum_{(i,j) \in \mathcal{C}} \Delta J_{ij}(a \to b, s_j). \tag{19}$$

Whereas in fully connected DCA this equation contains $L - 1$ $J$-based terms for each position $i$ (which scale with MSA length), the number of such terms is here reduced to the number of connections of $i$.

In our StructureDCA model, the sparsity $\mathcal{C}$ of the DCA model is defined by the residue–residue contacts extracted from the 3D structure of the protein (note, however, that we also provide users with full control over the sparsity). Based on the idea that residues in contact coevolve [10], the goal is to remove as many distant couplings as possible, which may introduce noise in the model, while retaining the most informative couplings between interacting residues. For that reason, we sometimes refer to the connection graph $\mathcal{C}$ as the contact map or contact matrix. We provide more technical details about the derivation of $\mathcal{C}$ and our sparse DCA PLM-based solver in Supplementary Section 2.

#### 1.8 Position and pairwise reweighting

Given a sparse DCA model with already inferred coefficients $\{h, J\}$, we can modify the equations for the probability of a sequence (Eq. 10), its energy (Eq. 9), and its energy change upon mutation (Eq. 12) by reweighting the relative contribution of each position and each pair of positions. For this purpose, we introduce positional weights $w_i^h$ and pairwise weights $w_{ij}^J$ (not to be confused with the sequence weights $w^n$). Using these coefficients, the energy of the model can be rewritten as:

$$E(s) = -\left( \sum_{i=1}^{L} w_i^h h_i(s_i) + \sum_{i<j \mid (i,j) \in \mathcal{C}} w_{ij}^J J_{ij}(s_i, s_j) \right). \tag{20}$$

with other DCA-derived quantities modified in a similar manner.

These weights aim to introduce an external source of information in addition to the DCA model. It is important to note that, in contrast to the sparsity modification, the introduction of weights is no longer strictly consistent with the statistical physics formalism of a generalized Potts model. Indeed, while a sparse DCA model still defines a probability distribution on the MSA that maximizes the likelihood of the observed sequences (under specific sparsity constraints), the weighted DCA energy and probability values are no longer guaranteed to correspond to a maximum-likelihood solution for the observed sequences. For this reason, the reweighting step should not be viewed as a modification of the underlying DCA model itself, but rather as an adjustment of the DCA-derived quantities that incorporates additional external information.

We showed in a previous study [4] that adding relative solvent accessibility (RSA)-based weights can substantially improve the performance of both profile and epistatic models (including DCA-based models) in predicting effects of mutations. However, in that case, the weights were applied only as positional multipliers after prediction and were not incorporated into the model itself. This made the approach suitable only for single-site mutations.

Here we generalizes this idea to fully integrated it into the DCA formalism. More in detail, we defined the positional and pairwise weights $w_i^h$ and $w_{ij}^J$ (as used in Eq. 20) as a function of the RSA of the corresponding residue(s) in the 3D structure. For single positions, the weight $w_i^h$ is defined as the complement of the RSA at position $i$,

$$w_i^h = 1 - \frac{\min(\mathrm{RSA}_i,\, 100)}{100}. \tag{21}$$

For pairs of positions, the weight $w_{ij}^J$ is defined as the average of the RSA complements at $i$ and $j$,

$$w_{ij}^J = \frac{w_i^h + w_j^h}{2} = \frac{\left(1 - \frac{\min(\mathrm{RSA}_i, 100)}{100}\right) + \left(1 - \frac{\min(\mathrm{RSA}_j, 100)}{100}\right)}{2}. \tag{22}$$

When these RSA-based weights are used, we refer to the model as StructureDCA[RSA]. Further technical details on the derivation of the RSA-based weights are provided in the Supplementary Section 2.

##### 1.9 Positional mutational tolerance

Among other measures, a (sparse) DCA model of a protein family can be used to compute the mutational tolerance [11] at each position of the MSA, which represents how permissive a given position is to amino-acid substitutions. As in Eq. 14, given a DCA model, we define the relative probability of observing amino acid $a$ at position $i$ given the background sequence $s$ as

$$P(i, a \mid s) = \frac{e^{-\Delta E(i, s_i \to a \mid s)}}{\sum_{b=1}^{q} e^{-\Delta E(i, s_i \to b \mid s)}}. \tag{23}$$

Note that this probability is independent of the wild-type amino acid $s_i$ at the target position, since it is normalized over all possible amino acid states $b = 1, \ldots, q$. Using the positional relative amino acid probabilities, we define the mutational tolerance at position $i$ given a background sequence $s$ as the Shannon entropy at this position,

$$MT(i \mid s) = -\sum_{a=1}^{q} P(i, a \mid s) \log_2 P(i, a \mid s). \tag{24}$$

By construction, $MT$ is zero when only one amino acid is allowed at this position and is maximal ($\log_2(q)$) when all amino acids are equally probable. Since mutational tolerance is derived from positional probabilities, which are independent of the wild-type amino acid $s_i$ at the target position, it is itself independent of $s_i$. This property is important, as it allows to compare mutational tolerances at position $i$ between two different background sequences $s$ and $s'$, even if they carry a different amino acid at this position.

In this work, we compute positional probabilities and mutational tolerances by considering only the 20 standard amino acids as possible states, excluding the gap state.

### 2 StructureDCA implementation

The StructureDCA implementation consists of the following steps, which we briefly summarize here and describe in detail in the next sections.

- **Structure processing** (Section 2.1). StructureDCA parses the provided structure file and extracts the following features: the amino acid sequence of the target chain(s), the RSA of each residue, the pLDDT or the B-factor (depending on whether the structure is experimental or predicted by AlphaFold or a similar method), and the residue–residue distance matrix.

- **MSA processing** (Section 2.2). StructureDCA parses the provided MSA file. It extracts the target amino acid sequence (defined as the first sequence of the MSA) and the remaining sequences. By default, it also removes redundant sequences and sequences that are too distant from the target sequence.

- **3D structure and MSA alignment** (Section 2.3). The aligner performs a pairwise alignment between the target sequence extracted from the MSA and the one extracted from the 3D structure in order to correctly map structural features to MSA positions or pairs of positions, even when the structure and MSA do not perfectly match. The aligner also extrapolates structural features to residues present in the MSA target sequence but missing in the structure.

- **Sparse DCA solver** (Section 2.4). Given a connection graph $\mathcal{C}$ (by default obtained from the contact map of the 3D structure) and the preprocessed MSA, the DCA coefficients $\{h, J\}$ are inferred using the sparse pseudo-likelihood maximization solver (implemented in C++ for performance reasons).

- **Structure-informed Direct Coupling Analysis** (Section 2.5). We explain how information from the corresponding 3D structure is exploited in the StructureDCA model, namely by using the contact map to define the sparsity of the DCA model and the RSA of residues to optionally define positional and pairwise weights.

#### 2.1 Structure processing

Protein 3D structures are parsed from a `.pdb` file. If the file contains multiple models, only the first model is considered and all others are ignored. As in RSALOR [12], the amino acid sequence of the target chain(s) is extracted, and non-standard amino acids are mapped to their corresponding standard amino acids using our curated mapping `AminoAcid._NON_STANDARD_AAS`. The mapping data is available at https://github.com/MatsveiTsishyn/NonStandardAminoAcidMapping and was established using data from the Protein Data Bank in Europe [13].

As we did in [12], StructureDCA computes the RSA using the Biopython module `Bio.PDB.SASA` [14] based on the Shrake and Rupley algorithm [15], which samples spheres of a given radius (simulating solvent molecules) to "probe" the surface of the molecule. The algorithm computes the solvent accessible surface area (SASA), which is then normalized to RSA (in %) by dividing the SASA of the residue in its structure by the SASA of the same residue in an extended tripeptide Gly-X-Gly conformation [16]. We use the empirically computed scale from [17] to normalize SASA to RSA values.

Values of pLDDT or B-factor (depending on whether the structure is experimentally solved or an AlphaFold (or similar) prediction) are extracted and assigned to each residue.

The distance matrix is computed only between residues of the target chain(s) and is accelerated using the Python package NumPy [18]. The distance is defined as the minimal atom-to-atom distance between two residues. By default, hydrogen atoms are excluded, as their presence depends on the origin of the input structure. Backbone atoms (except the $C_\alpha$ atom of glycine) are also excluded to avoid giving too much importance to the "diagonal contacts" arising from residues that are close in the protein sequence.

This behavior can be overwritten. Note that if these parameters are modified, the interpretation of the distance cutoff used to compute the contact map changes, as more atoms will be included in the distance calculation.

If multiple chains are specified as targets, the amino acid target sequence extracted from the 3D structure is the concatenated sequence of all these chains. In this case, StructureDCA expects that the MSA is a concatenated-MSA of the same sequences in the same order (otherwise the mapping of the 3D structure to the MSA might be wrong).

If the provided 3D structure represents a biologically meaningful homo-oligomeric complex, StructureDCA can also integrate inter-chain residue-residue interactions to compute the contact and distance matrices. For instance, if the 3D structure is a homo-dimer of chains `A` and `B`, the distance between residue 5 and 15 is computed as the minimum distance between $(A5, A15)$, $(A5, B15)$, $(B5, A15)$ and $(B5, B15)$.

Finally, the residue-residue contact matrix $C$ (often called the contact map) is computed from the distance matrix $D$ using the distance cutoff $d_0$,

$$C[i,j] = \delta_{D[i,j]<=d_0}, \tag{25}$$

where $\delta$ is the Kronecker delta.

Optionally, by setting `use_contacts_plddt_filter=True`, users can remove residue-residue contacts in poorly resolved ("spaghetti-like") regions by filtering out those where at least one residue has a low pLDDT score. However, contacts between such low-confidence residues may still be retained if the residues are close in sequence. For instance, if we set `contacts_plddt_cutoff=70` and `contacts_plddt_keep_window=2`, a residue $i$ with a poor pLDDT (e.g., 25) is only considered to be in contact with residues $i-2$, $i-1$, $i+1$ and $i+2$ provided that the distance cutoff criterion of Eq. 25 is satisfied.

#### 2.2 MSA processing

Input MSAs can be provided in `.fasta`, `.a2m`, or `.a3m` formats. They can be compressed as a `.gz` file.

StructureDCA considers the first sequence in the MSA as the target sequence of the model. This first sequence cannot contain any gaps (`-`) or non-standard one-letter codes like `B` or `X`, since it will be used to align positions in the MSA with residues in the 3D structure. However, users can choose to ignore the first sequence in the inference of the DCA coefficients if the model is required not to be biased towards this specific sequence. Insertions in the following sequences (relative to the target sequence) in the .a3m MSA format are ignored. All non-standard one-letter amino acid codes in these sequences are replaced with gaps (`-`).

StructureDCA removes duplicate sequences (exact matches between two sequences in the MSA). Note that it can happen that two full sequences are different but that their respective sequences restricted to the MSA are exactly the same.

StructureDCA also removes sequences from the "twilight zone" [19], i.e., sequences that are too distant from the target sequence based on a sequence identity criterion (by default 0.25).

#### 2.3 3D structure and MSA alignment

It is common for the target sequence in the MSA and the sequence extracted from the protein 3D structure to differ slightly. For example, many experimental protein structures from the Protein Data Bank (PDB) [20] contain missing residues, particularly at the termini of the protein chain. Additionally, residue numbering in PDB 3D structures often does not match the sequential indexing used in FASTA sequences. Another example of such a mismatch is when the provided MSA and 3D structure represent slightly different segments of a larger protein (as is the case for some proteins in ProteinGym [21]). Finally, a user might wish to evaluate the mutational landscape of a protein using the StructureDCA score derived from the structure of a homologous template protein that differs slightly from the target protein, assuming the overall secondary and tertiary structures are conserved. This scenario is reasonable, as the contact maps and RSA values tend to remain similar across homologous proteins with similar function.

All these scenarios make the manual process of mapping structural features to positions in the MSA tedious. StructureDCA automates this process by performing a pairwise alignment between the MSA target sequence and the sequence extracted from the protein 3D structure using the Biopython module `Bio.Align.PairwiseAligner` [14] with adjusted alignment parameters.

Once the alignment is done, structural features are mapped to MSA positions: RSA and pLDDT are expressed as vectors of length $L$, and the distance matrix is expressed as a matrix of size $L \times L$, where $L$ is the length of the target sequence in the MSA. Values of RSA, pLDDT, and distances for residues that are missing in the 3D structure are extrapolated based on the values of neighboring residues. The extrapolation is done with the idea that structurally missing residues are under-considered (or even ignored) rather than over-considered. In practice, their pLDDT is set to 0, their RSA rapidly increases to the maximal value of 100, and their distances to all other residues are rapidly incremented. Exact implementation is not described here; please refer to the source code for the `StructureSequenceAlignment` object. These extrapolations are implemented as a fail-safe option when a few residues are missing or when a small tail of the protein is missing. It is in no way a 3D structure prediction feature; therefore, the user should rely on 3D structure prediction models if a significant portion of the protein is not structurally resolved.

##### 2.4 Sparse DCA solver

Our sparse pseudo-likelihood maximization solver is implemented in C++ and supports multi-threading. Given an MSA and a connection graph $\mathcal{C}$, the solver infers sparse DCA coefficients $\{h, J\}$. The $J_{ij}$ coefficient matrices corresponding to a non-connected pair $\{i, j\}$ are never initialized. Since in StructureDCA $J$ is often expected to be highly sparse, it is stored using a memory-efficient sparse data structure, allowing for a substantial reduction in RAM usage.

In the first step, the solver computes weights for each sequence in the MSA as in Eq. 4 (with $\tau = 0.80$ by default). Next, the parameters $\{h, J\}$ of the Potts model are initialized to those of an independent-site Potts model with a zero-sum gauge, i.e.

$$\begin{cases} J = 0 \\ h = h^* = \log f_i(a) - \sum_{b=1}^{q} \log f_i(b) \end{cases} \tag{26}$$

where $f_i$ are the single-site amino acid frequencies at position $i$ (regularized and weighted as in Eq. 7). Note that this is the only moment in which StructureDCA uses the frequency-based pseudocount (parameter `theta_regularization`); its impact on the inferred coefficients is therefore very limited or even unnoticeable. This hot initialization helps to perform a slightly faster and more stable convergence. From this initial state, the parameters are then optimized to maximize the $L2$-regularized pseudo-likelihood $\mathcal{PL}^{L2,w}$ (Eq. 17), using gradient descent with the L-BFGS quasi-Newton scheme [22], as classically done in PLM-based DCA implementations [9, 23, 24]. Finally, the gauge is set to zero-sum [8].

For numerical stability and to avoid a scaling of the gradient with the number of effective sequences $N_{\text{eff}}$ (Eq. 5), the gradient is computed on the $L2$-regularized pseudo-likelihood $\mathcal{PL}^{L2,w}$ divided by $N_{\text{eff}}$. Note that, if the $L2$-regularization factors $\lambda_h$ and $\lambda_J$ are kept constant with respect to $N_{\text{eff}}$, they provide stronger regularization for low-depth MSAs, preventing overfitting to specific properties observed in a small number of sequences. With this formulation, the $L2$-regularization terms will therefore become negligible for larger $N_{\text{eff}}$ values. However, it may also be useful to maintain a certain amount of "asymptotic" regularization even when $N_{\text{eff}}$ becomes very large. Indeed, sequences in a given MSA will always contain some sampling bias, and there are still regions of the possible protein sequence space that remain unexplored by evolution. Moreover, this formulation also prevents possibly diverging values for the DCA coefficients. For that reason, the effective $L2$-regularization factors $\lambda_h$ and $\lambda_J$ used by StructureDCA are asymptotically corrected as

$$\lambda_{h/J} = \lambda^0_{h/J}\,(1 + N_{\text{eff}}\lambda^{\infty}) \tag{27}$$

where $\lambda^0_h$ and $\lambda^0_J$ are the initial factors, and $\lambda^{\infty}$ is the asymptotic correction. By default, we set $\lambda^0_h = \lambda^0_J = 1$ (parameters `lambda_h` and `lambda_J`) and $\lambda^{\infty} = \frac{1}{1000}$ (parameter `lambda_asymptotic`), corresponding to an $N_{\text{eff}}$ value of 1000, which represents a relatively deep MSA around the median values among proteins from ProteinGym. This means that even if $N_{\text{eff}} \to \infty$, the relative magnitude of the $L2$-regularization term remains comparable to that obtained with a regularization factor $\lambda^0_{h/J}$ for an MSA with $N_{\text{eff}} = 1000$.

#### 2.5 Structure-informed DCA

We put all the pieces together and define the StructureDCA model and its RSA-reweighted variant, StructureDCA[RSA]. We explain here how exactly StructureDCA leverages information from the corresponding 3D structure to adapt its underlying DCA model.

StructureDCA uses the contact map $C$ derived from the 3D structure (Eq. 25; optionally including the pLDDT criterion) to define the sparsity of the underlying DCA model (namely, the connection graph $\mathcal{C}$; see Supplementary Section 1.7). The distance threshold $d_0$ is set by default to 8Å (parameter `distance_cutoff`). In addition, we provide the user with full control over the sparsity.

As shown in the Supplementary Section 1.8, StructureDCA can be reweighted by introducing positional weights $w^h_i$ and pairwise weights $w^J_{ij}$ in the evolutionary energy. In the StructureDCA[RSA] model, these weights are functions of RSA (see Eqs. 21 and 22). As discussed in the main text, this model is better suited for capturing the thermodynamic stability properties of proteins, although it may be less relevant for functional or fitness-related properties. To apply these RSA-based weights when evaluating the energy of a sequence, $E$, or the energy change upon mutation, $\Delta E$, users must set the parameter `reweight_by_rsa=True`.

Note that we also provide users with full control over the weights used. This means that users can customize both positional and pairwise weights; for instance, one may assign to $w^h_i$ a measure of the closeness of residue $i$ to a functional site, and to $w^J_{ij}$ the average closeness of residues $i$ and $j$ with respect to that site.

#### 2.6 StructureDCA usage

StructureDCA is distributed as a Python package available on PyPI and can be installed with the command `pip install structuredca`. It is free and open source under the MIT license, and the source code is available on its GitHub repository.

Its sparse DCA solver is implemented in C++ for performance, and we provide a user-friendly and fully customizable Python interface as well as a command-line interface. We also provide Jupyter notebook tutorials in the GitHub repository, illustrating different usage scenarios. For instance, we illustrate how users can change default model sparsity and positional and pairwise RSA weights.

StructureDCA can also be executed through a graphical interface, via its Google Colab Notebook, allowing its usage without writing code or installing software. The Colab interface enables users to upload or fetch 3D structures (from AlphaFold-DB [25] or the PDB [20]) and MSAs (uploaded, retrieved, or generated via the MMseqs2 API [26]). When multiple chains are present in the structure, the available chains are listed, and alignment feedback is provided to ensure consistency between the selected chain and the MSA target sequence. Finally, it provides graphical visualizations of the outputs.

#### 3 Evaluation metrics and datasets

As main evaluation metric, we use Spearman's rank correlation coefficient (which we refer to as the "Spearman correlation" or $\rho$) between experimental DMS measurements and predicted values. First, Spearman correlations are computed for each individual DMS experiment and then averaged across all experiments within the DMS collection (ProteinGym, MegaScale and HumanDomains). The Spearman correlation measures the Pearson correlation between the ranks of two sets of values and is therefore well suited for comparing quantities that may have different units or exhibit non-linear relationships. This is the case for both computational predictions and experimental measurements in ProteinGym. For consistency, we use the Spearman correlation as the main metric in all evaluations. For completeness, we also report the Pearson correlation for HumanDomains and MegaScale.

As each DMS collection has its own specificities and requires its own corrections, we use different aggregation schemes for individual Spearman correlations. We therefore clarify here what we mean by "average Spearman correlation" for each of the DMS collections used.

- **ProteinGym:** We aggregate Spearman correlations following the procedure of the original benchmark [21]. The 217 datasets are first grouped by DMS type ("stability", "binding", "expression", "activity", and "fitness"). For each DMS type, the average Spearman correlation is computed by reweighting the contribution of each dataset by clustering together datasets with the same target protein. Then, the overall Spearman correlation on the ProteinGym benchmark is computed as the average across the five DMS type, such that each category contributes equally, regardless of the number of datasets it contains.
- **MegaScale:** To account for the presence of highly similar target proteins in MegaScale (sometimes differing by only a single mutation), we reweight the dataset contributions by clustering proteins that share at least 80% sequence identity (using the CD-HIT software for clustering [27]). When comparing StructureDCA with other computational models from the benchmark in [4], we limited the evaluation to datasets and mutations included in that benchmark, so that all models are assessed on identical data. The final subset used for this evaluation contains around $120,000$ mutations from 115 individual DMS experiments and includes only single-site mutations. This subset corresponds to the intersection between our data and the benchmark from [4].
- **HumanDomains:** In this case, we simply compute the average Spearman correlation across all DMS experiments. When comparing StructureDCA with other computational models from the benchmark in [28], we restricted the evaluation to datasets and mutations for which the benchmark provides predictions for all evaluated models, ensuring that all models are assessed on the same data. After this filtering, any dataset containing fewer than 100 mutations was discarded to avoid calculating correlations on very small samples. The final subset used for this evaluation contains around $400,000$ mutations from 392 individual DMS experiments. This subset corresponds to the intersection between our data and the benchmark from [28].

### 4 StructureDCA computational time complexity and efficiency

We quantify how StructureDCA reduces the size and computational cost compared to a standard fully connected DCA model. As the number of $J$ coefficients increases proportionally to the square of the sequence length $L$, the number of DCA parameters can become extremely large. For instance, even a relatively short protein of length $L = 200$ requires nearly ten million coupling coefficients. Alternatively, the problem can be reformulated in terms of the average number of couplings per position in the MSA. In a standard fully connected DCA model, this number scales linearly with the sequence length $L$, since couplings with all other positions must be considered. As a consequence, fully connected DCA models involve a large number of terms to evaluate the effect of a single mutation (Eq. 12), which can introduce noise in predictions. In contrast, for StructureDCA (for a fixed distance cutoff $d_0$), the average number of couplings $C$ per position remains limited, as only a small number of residues can fit within a spherical volume of radius $d_0$.

In Fig. 2d of the main publication, we show how the average number of couplings per position $C$ scales with protein length $L$ for both standard DCA and StructureDCA, using all proteins from the ProteinGym dataset. As $L$ increases, the ratio between $C$ in fully connected DCA and in StructureDCA grows rapidly. The average number of couplings per position in StructureDCA (with $d_0 = 8$ Å) remains approximately constant and bounded between 14 and 18. In terms of model size (number of DCA coefficients), StructureDCA achieves a reduction of roughly 4-fold for small domains with $L = 50$, about 30-fold for proteins of length $L = 500$, and up to 360-fold for the largest proteins in ProteinGym.

In terms of computational cost, standard pseudo-likelihood maximization DCA methods scale as $\mathcal{O}(NL^2)$, where $N$ is the number of sequences in the MSA and $L$ its length. At each gradient-descent iteration, all $N$ sequences must be processed and all $\mathcal{O}(L^2)$ coupling coefficients updated. In contrast, StructureDCA reduces this complexity to $\mathcal{O}(NL)$, as the number of coupling coefficients now scales linearly with $L$.

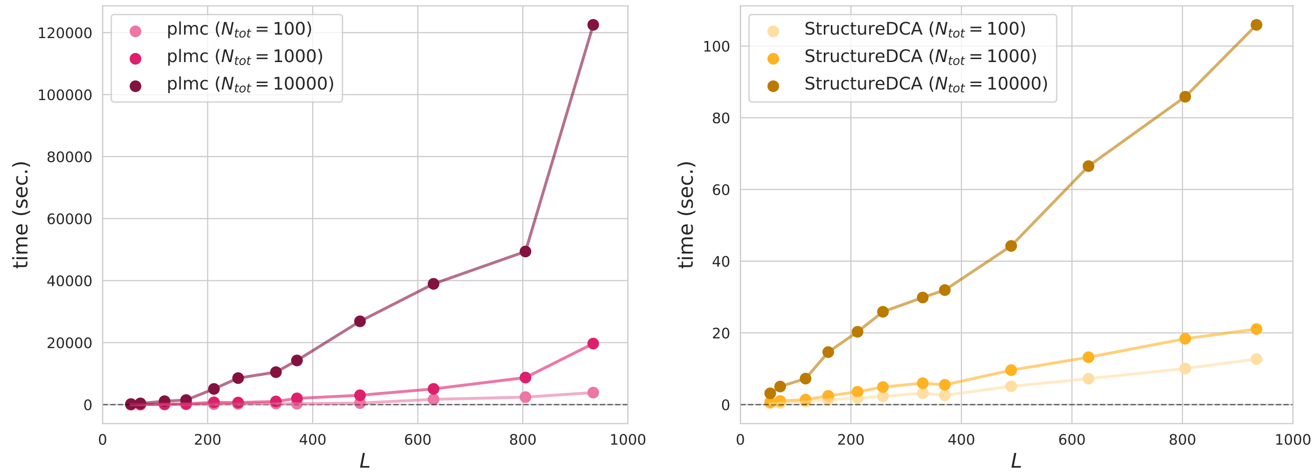

**Figure S1: Computational times of StructureDCA vs. standard DCA (plmc).**
We compare the execution times of StructureDCA with those of plmc, the DCA solver used by EVmutation [23], which is also based on the pseudo-likelihood maximization method. plmc was executed with default parameters, and StructureDCA with default parameters (so using a distance cutoff of 8 Å) and with the sequence-identity filter disabled to avoid applying any pre-processing to the MSA. The evaluation was performed on a sample of proteins from ProteinGym (see Tab. S1). The sample was selected to span an MSA length range between $L = 50$ and $L = 1000$, and to contain at least 10000 distinct sequences in the MSA. To control for a single parameter at a time (the length or depth of the MSA), we subsampled each MSA to randomly retain $N_{\text{tot}} = 100$, $N_{\text{tot}} = 1000$, or $N_{\text{tot}} = 10000$ sequences.
StructureDCA shows up to a 1000-fold improvement in computational time for MSAs of length around $L = 1000$. This difference probably becomes even larger as $L$ increases.
Benchmarks were performed on a Linux machine using 8 CPUs with an AMD EPYC 7763 processor.

We performed an empirical comparison with plmc, the DCA solver underlying EVmutation [23] (see Fig. S1 and Tab. S1). Remarkably, the observed speed-up exceeds what would be expected from model-size reduction alone. Computational time is reduced by several dozen-fold for proteins of length $L = 50$, several hundred-fold around $L = 200$, and roughly 1000-fold for $L = 1000$, with even larger gains expected for longer sequences. For example, we evaluated all 2.5 million ProteinGym mutations (217 datasets) in only a few hours on a standard 8-CPU laptop, with per-dataset runtimes ranging from a few seconds to a few minutes.

For the B1 metallo-$\beta$-lactamase (MBL) family [11], we compared the computational time required by StructureDCA and Boltzmann machine DCA (bmDCA; `adabmDCA 2.0` [29]) known to be computationally expensive. A single bmDCA run on this MSA required approximately 13 hours on 8 CPUs, whereas StructureDCA completed in only 10 seconds, corresponding to a $\sim$ 4000-fold speedup. Even when bmDCA was executed on a high-performance GPU (NVIDIA A40), StructureDCA still achieved roughly a 200-fold speedup on a standard 8-CPU laptop.

**Table S1: Computational times of StructureDCA vs. standard DCA (plmc).**
This table lists the selected proteins from ProteinGym used for the computational time benchmark shown in Fig. S1. Execution times for plmc and StructureDCA with an MSA depth of $N_{\text{tot}} = 10000$ are reported in seconds, along with the ratio between the two runtimes.

| **Dataset** | **MSA length** | **plmc** (sec.) | **StructureDCA** (sec.) | **time ratio** |
|---|---|---|---|---|
| RAD_ANTMA_Tsuboyama_2023_2CJJ | 54 | 177 | 3 | 59.0 |
| CATR_CHLRE_Tsuboyama_2023_2AMI | 72 | 395 | 5 | 79.0 |
| PHOT_CHLRE_Chen_2023 | 118 | 1101 | 7 | 157.3 |
| UBC9_HUMAN_Weile_2017 | 159 | 1469 | 14 | 104.9 |
| ESTA_BACSU_Nutschel_2020 | 212 | 5057 | 20 | 252.8 |
| CASP3_HUMAN_Roychowdhury_2020 | 258 | 8550 | 25 | 342.0 |
| MTH3_HAEAE_RockahShmuel_2015 | 330 | 10468 | 29 | 361.0 |
| SERC_HUMAN_Xie_2023 | 370 | 14243 | 31 | 459.5 |
| CP2C9_HUMAN_Amorosi_2021_activity | 490 | 26867 | 44 | 610.6 |
| SC6A4_HUMAN_Young_2021 | 630 | 38952 | 66 | 590.2 |
| ACE2_HUMAN_Chan_2020 | 805 | 49389 | 85 | 581.0 |
| MSH2_HUMAN_Jia_2020 | 934 | 122504 | 105 | 1166.7 |

### 5 Results: Supplementary Tables and Figures

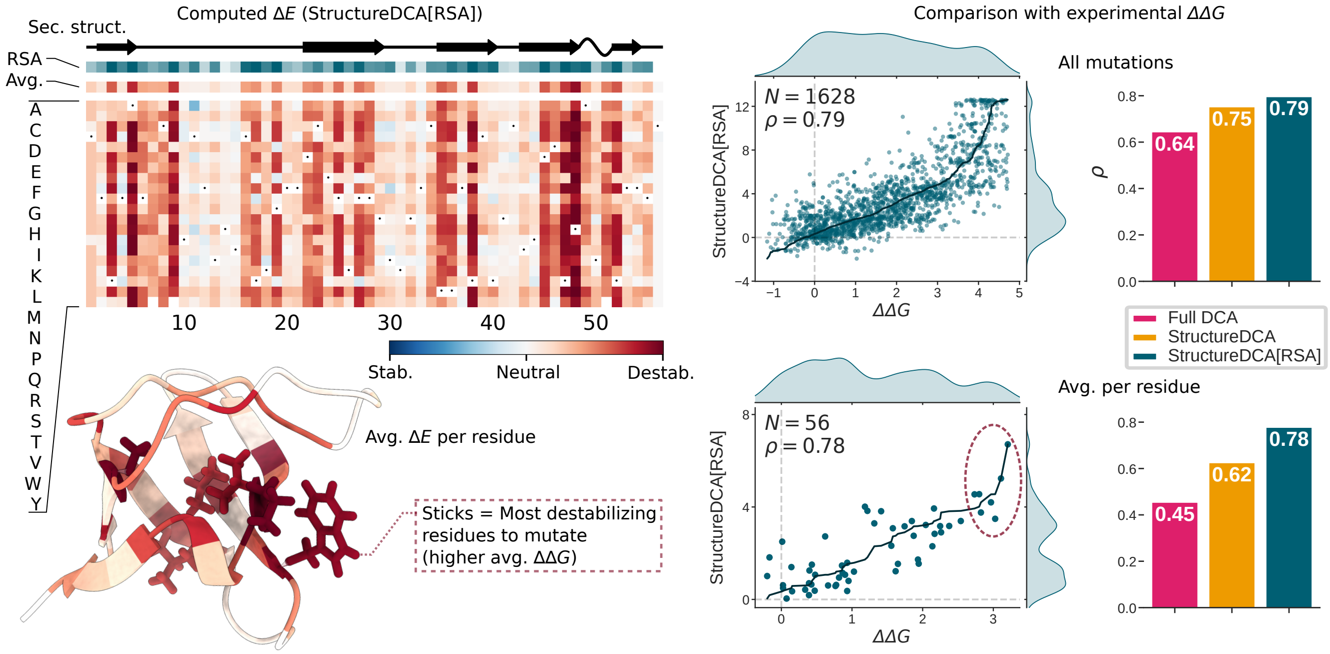

**Figure S2: StructureDCA and StructureDCA[RSA] output examples.**
As an illustration of our model, we show StructureDCA and StructureDCA[RSA] predictions for the stability of the SH3 domain of spectrin alpha (Gallus gallus) from the MegaScale dataset collection (referenced as `2cdt_A_7-62`). While standard DCA already performs well on this dataset ($\rho = 0.64$), it is largely outperformed by StructureDCA ($\rho = 0.75$) which is further improved by StructureDCA[RSA] ($\rho = 0.79$). These differences are even more pronounced when the results are averaged per residue, indicating that StructureDCA is able to correctly identify residues that are critical for protein stability.
On the left, we show a heatmap representation of StructureDCA[RSA], $\Delta E$ values for all single missense mutations and per-residue average values mapped onto the corresponding 3D structure. Residues that are the most sensitive to mutations for stability, as determined experimentally (circled in red in panel (b)), are shown in sticks. We observe that these residues are also predicted by StructureDCA[RSA] to be highly sensitive.
On the right, comparison of predictions for standard DCA (magenta), StructureDCA (orange) and StructureDCA[RSA] (blue) with experimentally measured stability changes upon mutation, $\Delta\Delta G$. The comparison is performed both on all mutations in the dataset (top; including single and double mutations) and on per-residue average values (bottom; averaging only over single-site mutations). Standard DCA was computed using our StructureDCA software by deactivating the structural filter on $J$ coefficients ($d_0 \to \infty$). StructureDCA[RSA] predictions are plotted against experimental $\Delta\Delta G$ values with respective marginal distributions. Spearman correlations ($\rho$) and number of entries ($N$) are are shown.

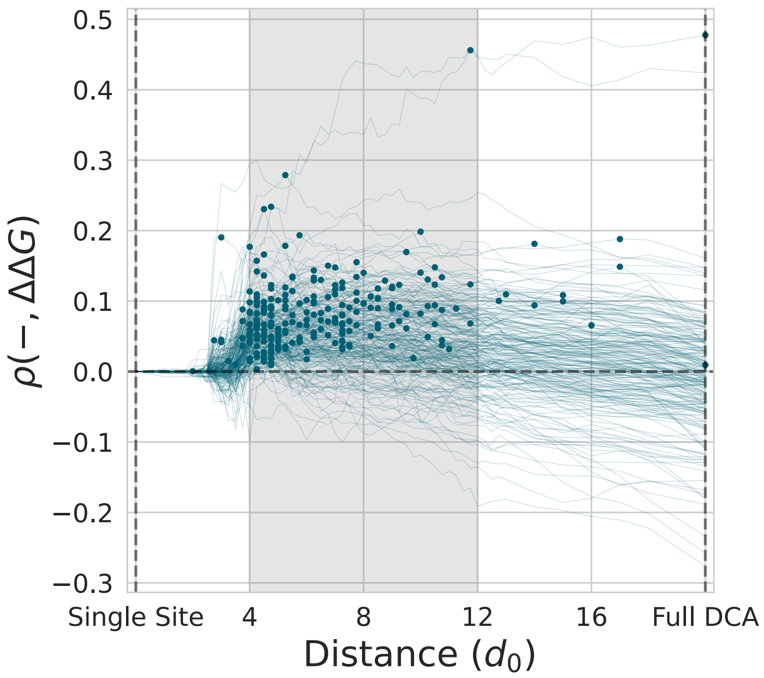

**Figure S3: StructureDCA distance spectrum on MegaScale.**
Spearman correlation curves between experimental stability measurements and StructureDCA predictions as a function of the distance cutoff $d_0$ (in Å) for each individual dataset in MegaScale. All curves were shifted such that the correlation at $d_0 = 0$ is zero, in order to highlight the impact of the distance cutoff instead of the overall performance trend on the dataset. Dots indicate the optimal point of each curve, i.e., the distance cutoff at which the Spearman correlation with experimental measurements is maximal. The leftmost value on the $x$-axis corresponds to the single-site model that considers no couplings ($d_0 = 0$). The rightmost values on the $x$-axis (placed at $x = 20$) correspond to the fully connected DCA model that includes all possible couplings ($d_0 \to \infty$).

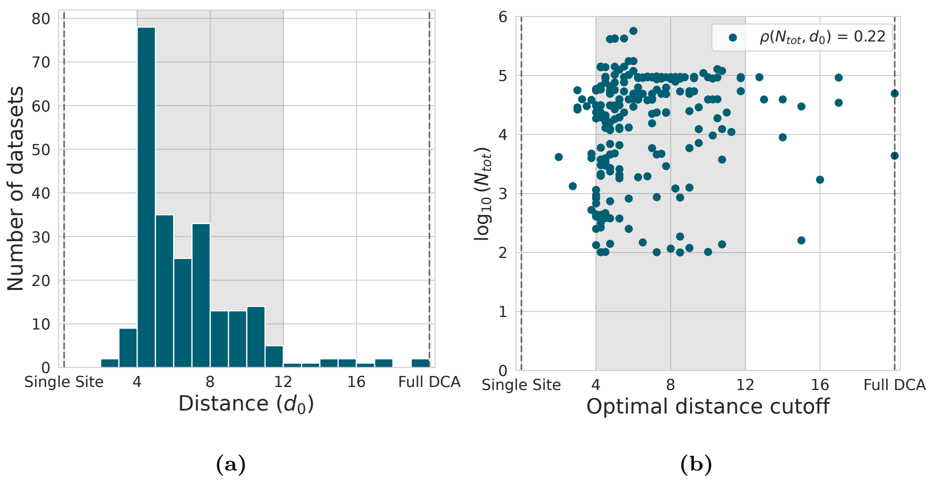

**Figure S4: StructureDCA optimal distances on MegaScale.**
For each individual dataset in MegaScale, we analyze the optimal distance cutoff $d_0$ (in Å), i.e., the distance cutoff at which the Spearman correlation with experimental measurements is maximal.
(a) Histogram of optimal distances.
(b) Comparison between the optimal distance and the number of distinct sequences in the corresponding MSA ($N_{\text{tot}}$, shown on a $\log_{10}$ scale). A slight positive correlation is observed between the two quantities (Spearman $\rho$ of 0.22).

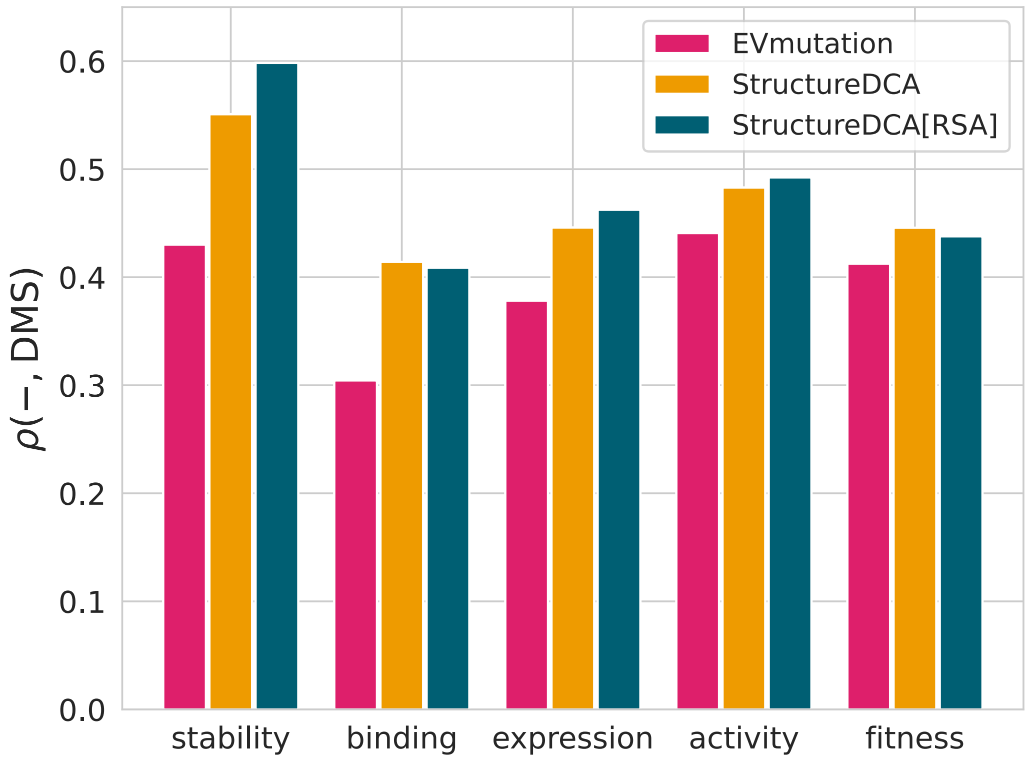

**Figure S5: Average Spearman correlation on ProteinGym by DMS assay type.**
Performance of EVmutation [23] (magenta), StructureDCA (orange), and StructureDCA[RSA] (blue) across stability, binding, expression, activity, and fitness assays. EVmutation is a standard fully connected DCA method based on pseudolikelihood maximization and serves as a baseline for comparison with StructureDCA.

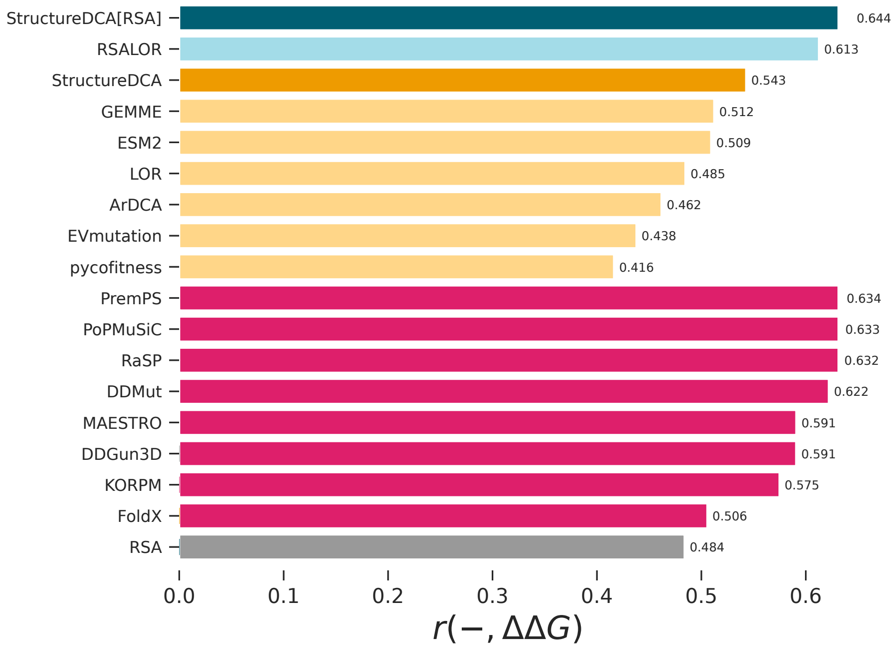

**(a)** MegaScale

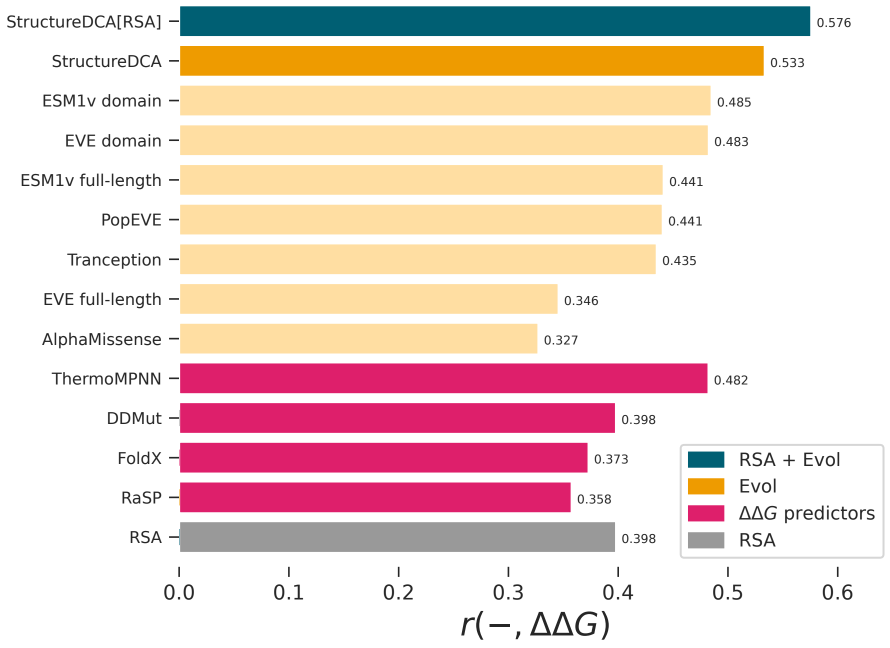

**(b)** HumanDomains

**Figure S6: Benchmark of StructureDCA, StructureDCA[RSA], and state-of-the-art methods for predicting mutational effects.**
Performance is evaluated as the average Pearson correlation ($r$) between predictions and experimental measurements.
(a) Performance on the MegaScale dataset collection for all methods benchmarked in [4], including classical structure-based $\Delta\Delta G$ predictors (magenta), evolutionary methods (orange), evolutionary methods augmented with RSA (blue), and RSA alone (grey).
(b) Performance on the HumanDomains dataset collection for methods benchmarked in [28], using the same color coding as in (a).
StructureDCA and StructureDCA[RSA] are highlighted in bright orange and bright blue, respectively.

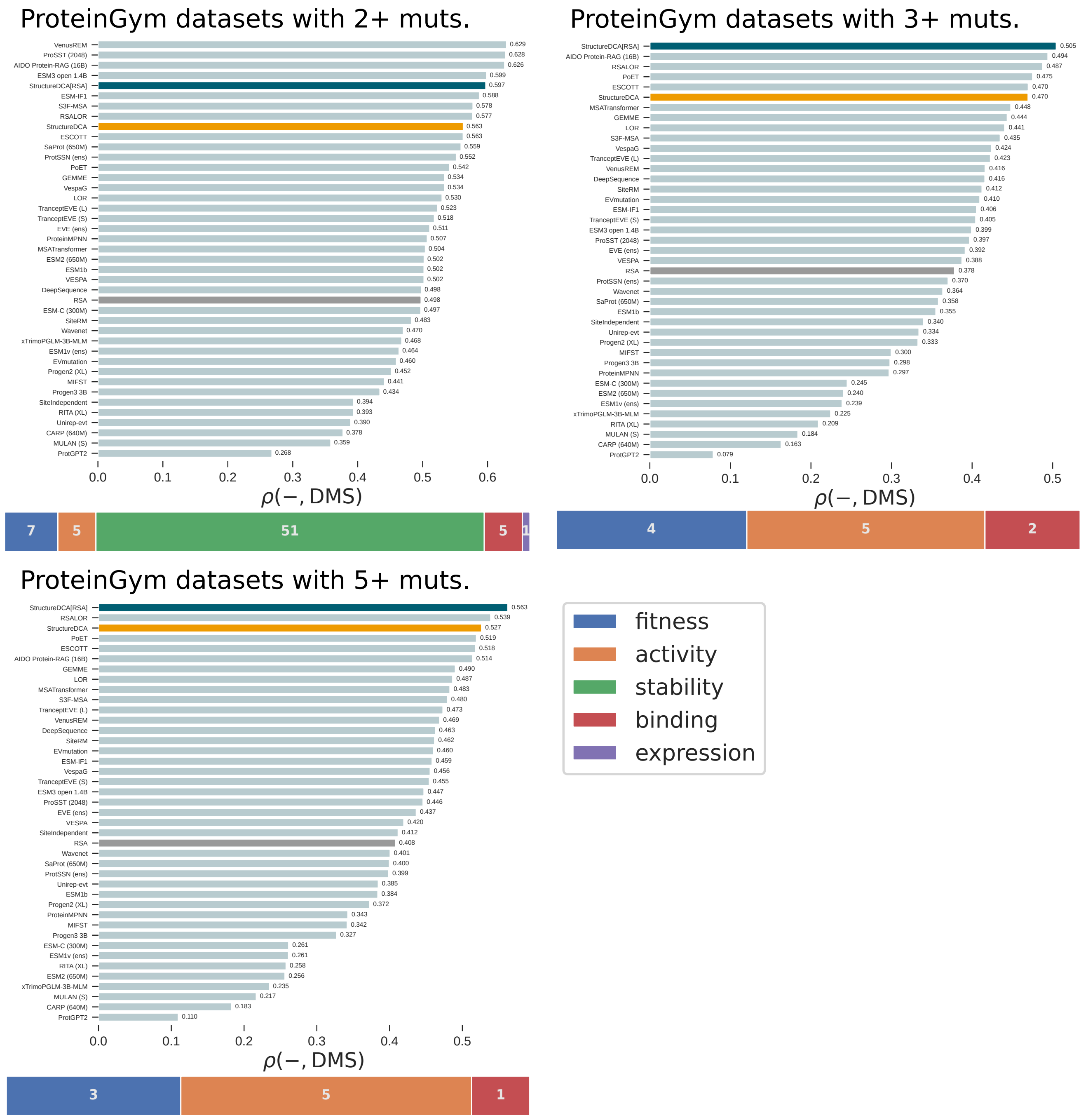

**Figure S7: Benchmark of StructureDCA, StructureDCA[RSA], and state-of-the-art methods for predicting mutational effects on ProteinGym datasets with higher-order mutations.**
Performance is measured as the average Spearman correlation. Methods from the original benchmark [21] are included, retaining only the best-performing model from each publication. Performances are evaluated on the 69 ProteinGym datasets containing multiple mutations with 2 or more sites mutated simultaneously, on the 11 datasets containing multiple mutations with 3 or more sites mutated simultaneously and on the 9 datasets containing multiple mutations with 5 or more sites mutated simultaneously. The case of 4 mutations is not shown as these are exactly the same ProteinGym datasets as for the 3 mutations. The legend indicates the number of datasets for each DMS type. StructureDCA, StructureDCA[RSA], and RSA are highlighted in bright orange, blue, and grey, respectively.

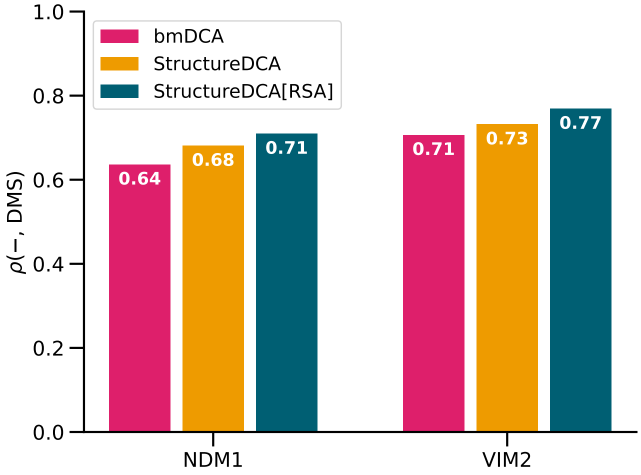

**(a)** Shared MBL family MSA

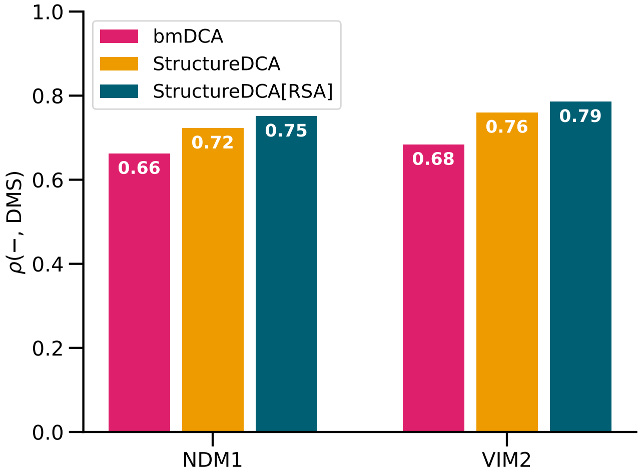

**(b)** Protein-specific MSAs (NDM1 and VIM2)

**Figure S8: Spearman correlations between DMS experimental values and DCA predictions for NDM1 and VIM2 metallo-$\beta$-lactamases (MBLs).**
DMS data for NDM1 was taken from [11] (selection with antibiotic ampicillin 'AMP' at 256 µg/mL, 37°C) and for VIM2 from [30] (ampicillin 'AMP' at 128 µg/mL, 37°C).
In addition to StructureDCA (orange) and StructureDCA[RSA] (blue), standard DCA predictions were computed using bmDCA (magenta) to reproduce the computational setup of [11]. bmDCA was run on GPU using `adabmDCA 2.0` [29] with parameter `-nsweeps 40` and default parameters otherwise. Reproduced results are very similar, though not identical, to aggregated values reported in [11], likely due to small differences in the MSA or software version. Only mutations at positions aligned to the MBL family MSA were considered. Because the NDM1 and VIM2 sequences aligned to the MBL family MSA contain some gaps, StructureDCA was run with parameter `exclude_gaps=False`.
(a) Using the MBL family MSA from [11] ($3,496$ sequences), which does not include NDM1 or VIM2, a single DCA/StructureDCA model was inferred and evaluated on both target proteins by varying only the background sequence. Used StructureDCA model incorporated both experimental structures by defining a contact if present in at least one structure and averaging RSA values across both structures. To use the provided MSA without any pre-processing, we disabled the sequence identity filter `min_seqid=None` for StructureDCA.
(b) Separate MSAs were generated for NDM1 and VIM2 following the usual procedure described in Methods. For consistency between the two MSA strategies, these MSAs were subsequently trimmed to match the same 222 positions of the MBL family MSA used in (a). StructureDCA was run with default parameters except `exclude_gaps=False`, and bmDCA with parameter `-nsweeps 40`. In this case, each StructureDCA prediction used the corresponding experimental structure.
Across both target proteins and both MSA strategies, StructureDCA consistently outperforms bmDCA, and StructureDCA[RSA] further improves the predictions.
As reflected by the overall improved performance of StructureDCA under strategy (b) compared to strategy (a), the carefully curated MBL family MSA is able to capture epistasis across distant homologs, while protein-specific MSAs appear to provide more accurate predictions of individual mutational landscapes.

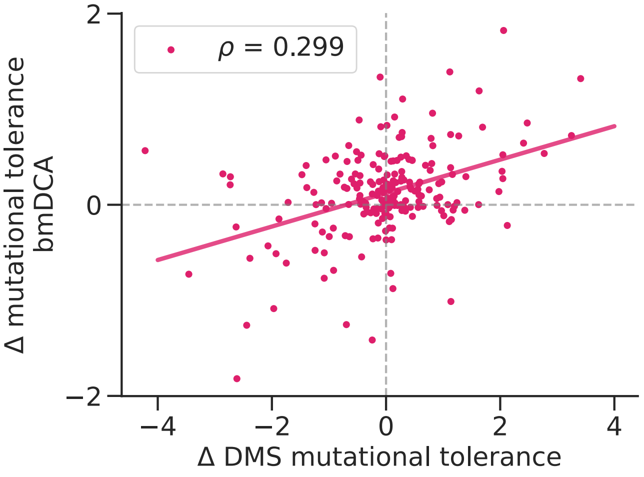

**(a)** bmDCA

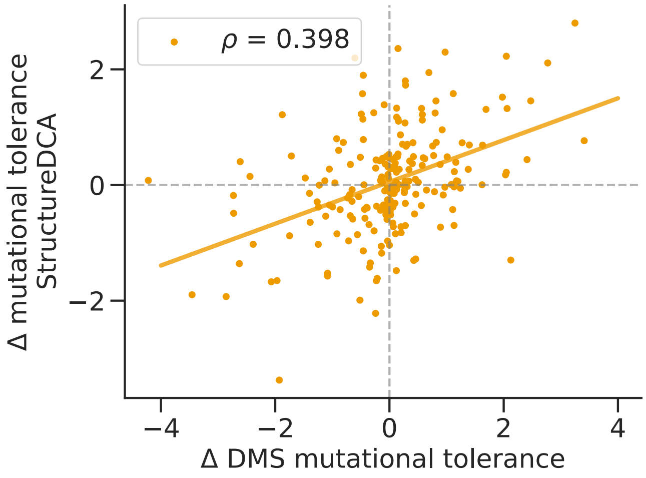

**(b)** StructureDCA

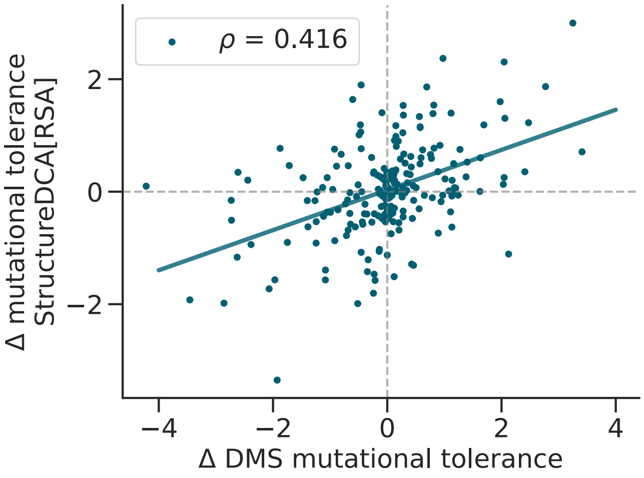

**(c)** StructureDCA[RSA]

**Figure S9: Comparison of mutational tolerance differences between NDM1 and VIM2 from DMS data and DCA predictions.**
For each position $i$ in the MBL family MSA, we compute the difference in mutational tolerance (Supplementary Section 1.9) between NDM1 and VIM2, $\Delta MT(i) = MT(i \mid \text{NDM1}) - MT(i \mid \text{VIM2})$.
DCA-based mutational tolerances are computed using bmDCA (magenta, standard DCA), StructureDCA (orange), and StructureDCA[RSA] (blue), all derived on the same MBL family MSA with parameters as in Fig. S8a. Since the same DCA/StructureDCA model is used for both proteins, the difference can only arise from the background sequence, thereby capturing epistatic effects. In contrast, a profile model such as LOR (or RSALOR) [4, 12], which is independent of the background sequence, would yield zero difference between the two proteins.
Experimental positional mutational tolerances are derived from raw DMS scores using Shannon entropy, following several normalization steps described in [11].
Lines represent linear regressions, and the corresponding Spearman correlations $\rho$ are shown.
All three DCA models capture the epistatic differences in mutational tolerance between NDM1 and VIM2. Consistent with previous results, StructureDCA agrees more with experimental data than bmDCA, with further improvement provided by StructureDCA[RSA].

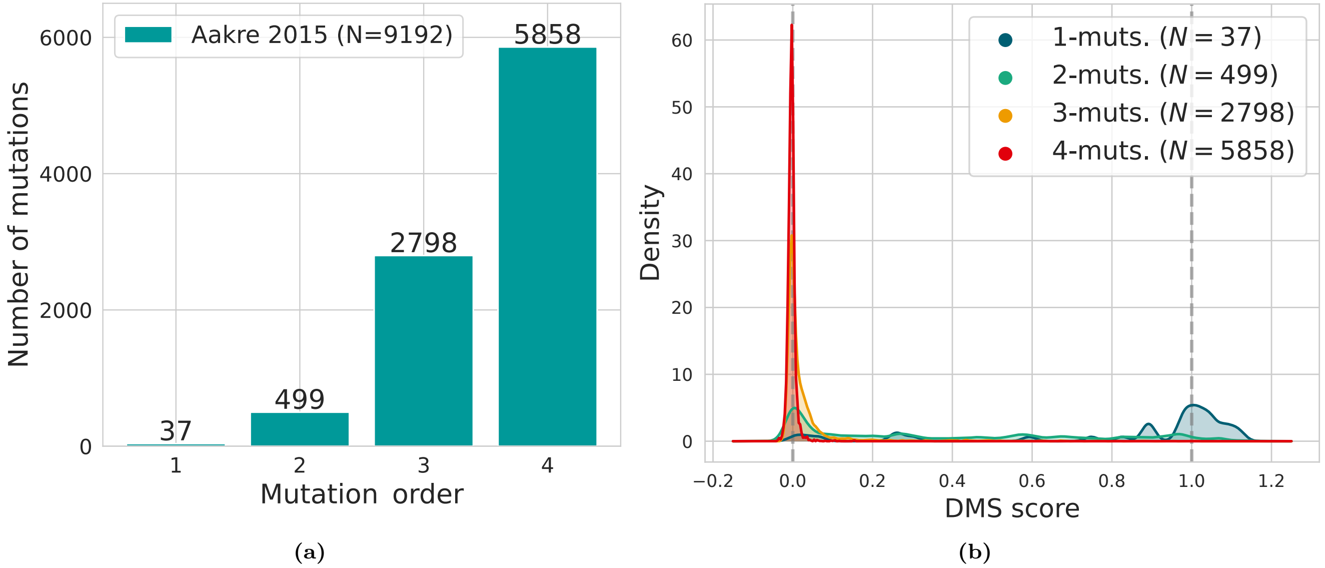

**Figure S10: DMS experiment on the ParD–ParE toxin–antitoxin complex [31].**
ParD–ParE is a toxin–antitoxin complex composed of the antitoxin ParD (UniProtKB `F7YBW8`) and its cognate toxin ParE (UniProtKB `F7YBW7`) in the prokaryotic species *Mesorhizobium opportunistum* (Taxon ID `536019`). This system provides a well-suited case study for coevolution in protein–protein interactions, as the primary evolutionary role of the antitoxin is to bind the toxin, imposing strong selective pressure on the interaction and on antitoxin fitness. In addition, the prokaryotic context limits the presence of paralogs, resulting in a clearer coevolutionary signal between the two proteins [32].
The mutagenesis experiment targets four sites on ParD (`L59`, `W60`, `D61`, and `K64`) and explores combinatorial mutations of orders one to four. The assay measures organismal fitness, which is strongly influenced by the ability of ParD to bind ParE.
(a) Number of mutations for orders 1, 2, 3, and 4. The number of mutations increases sharply with mutation order, with the majority of mutations corresponding to order 4.
(b) Distributions of DMS fitness scores for each mutation order. Mutations of order 4 exhibit fitness scores tightly clustered around zero, making correlation-based analyses less informative for this subset (for example, differences between fitness values of 0.01 and 0.02 are unlikely to be biologically meaningful in a scale from 0 to 1).

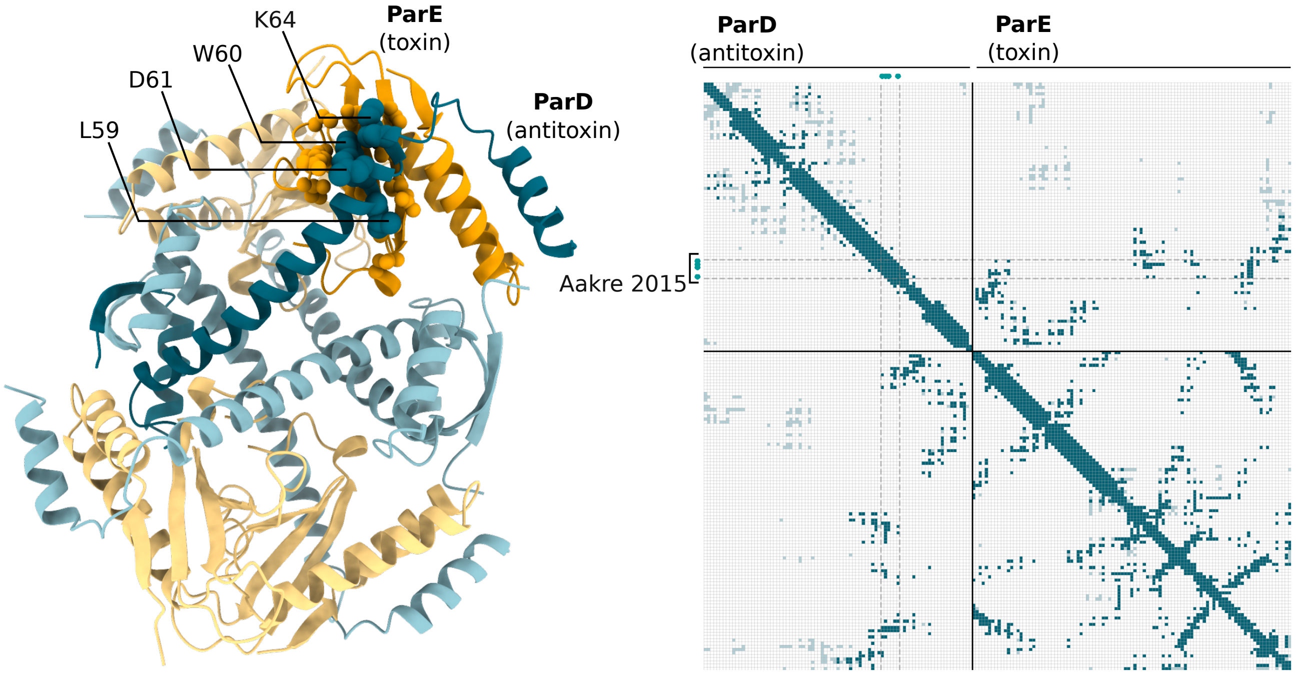

**Figure S11: Structural context of the four mutated residues in the ParD-ParE DMS dataset [31].**
Left: Experimental 3D structure of the ParD–ParE complex in its biological unit (PDB `5ceg` provided in the same publication [31]). As shown by the authors, one ParD antitoxin chain (blue) and one ParE toxin chain (orange) form a tightly interacting heterodimer, and four such heterodimers assemble into an octameric complex. For clarity, one heterodimeric subunit is highlighted using more saturated colors. The four target residues that are mutated (`L59`, `W60`, `D61`, and `K64`) on ParD are shown as blue spheres (displayed only once on the highlighted heterodimer).
Right: Joint ParD-ParE contact map computed by StructureDCA (with default distance threshold of 8Å). The four target residues are indicated by blue dots along the ParD axis. Contacts arising within the target heterodimeric ParD–ParE complex are shown as darker squares, while additional contacts introduced by the full octameric assembly are shown as lighter squares. As illustrated by both the 3D structure and the contact map, the four target residues participate only in sequence-local intra-chain contacts within the same $\alpha$-helix, which are expected to provide limited functional information about the ParD–ParE complex. In contrast, these residues are located at the ParD–ParE interface and exhibit multiple contacts with ParE.
For biological accuracy, StructureDCA computations were therefore performed using the contact map derived from the full octameric biological unit rather than from the heterodimer alone. While this choice may affect the global StructureDCA model, the evolutionary energy changes $\Delta E$ associated with the four target residues remain largely unchanged, as the octameric assembly introduces no additional contacts involving these positions.

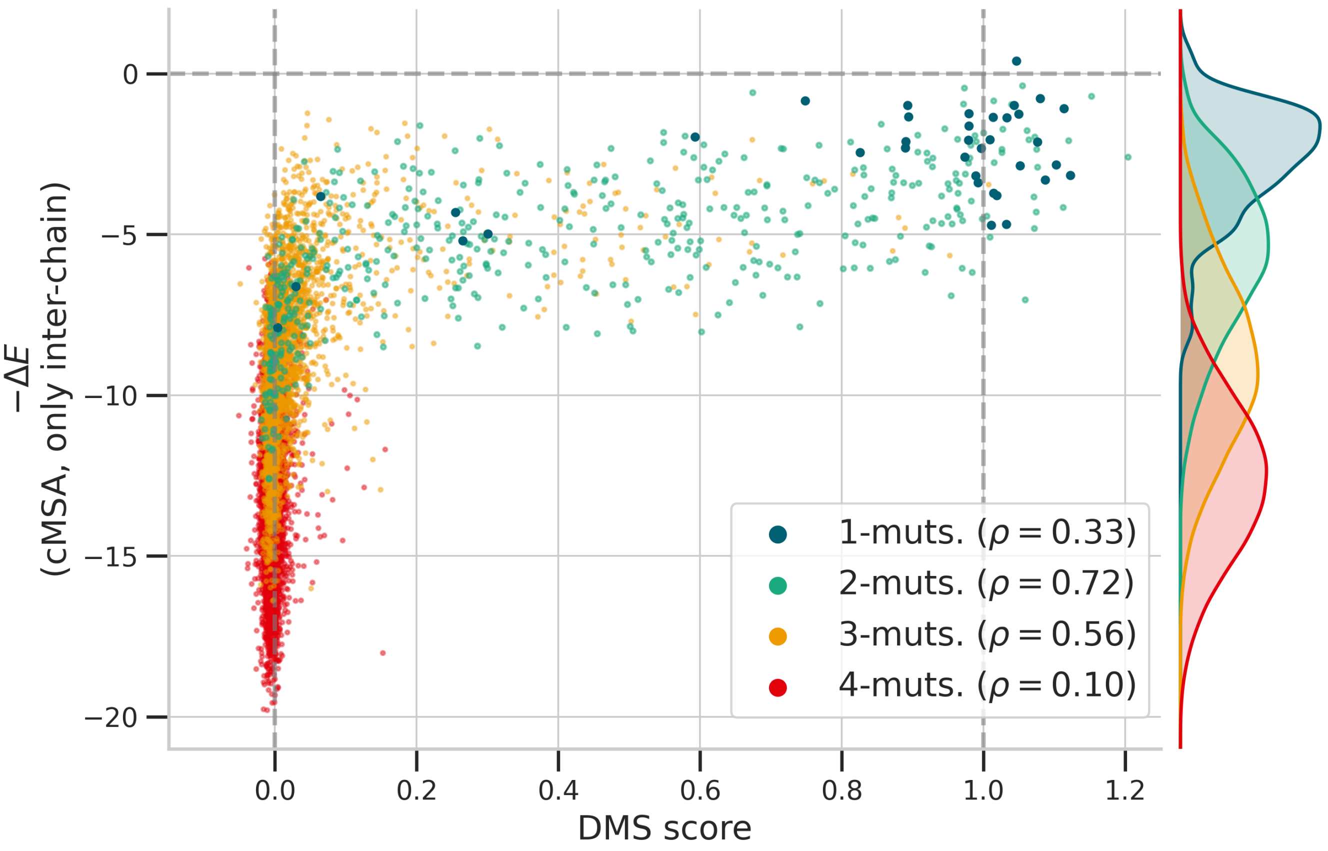

**Figure S12: Comparison between DMS fitness scores and StructureDCA predictions in the ParD-ParE DMS dataset [31].**
Scatter plot of experimental DMS fitness scores versus the negative StructureDCA evolutionary energy, $-\Delta E$, computed using the concatenated ParD–ParE multiple sequence alignment (cMSA) and considering only inter-chain couplings between ParD and ParE residues. The contact map is computed on the full octameric biological unit (experimental structure with PDB code `5ceg` [31]). Points are colored according to mutation order. On the right, the marginal distributions of $-\Delta E$ values are shown for each mutation order.
We observe that StructureDCA predictions show notable correlations with experimental values across all mutation orders, except for order 4, where experimental fitness scores are all clustered around zero (making correlations less informative for this subset). They are nonetheless correctly predicted by StructureDCA as predominantly deleterious.

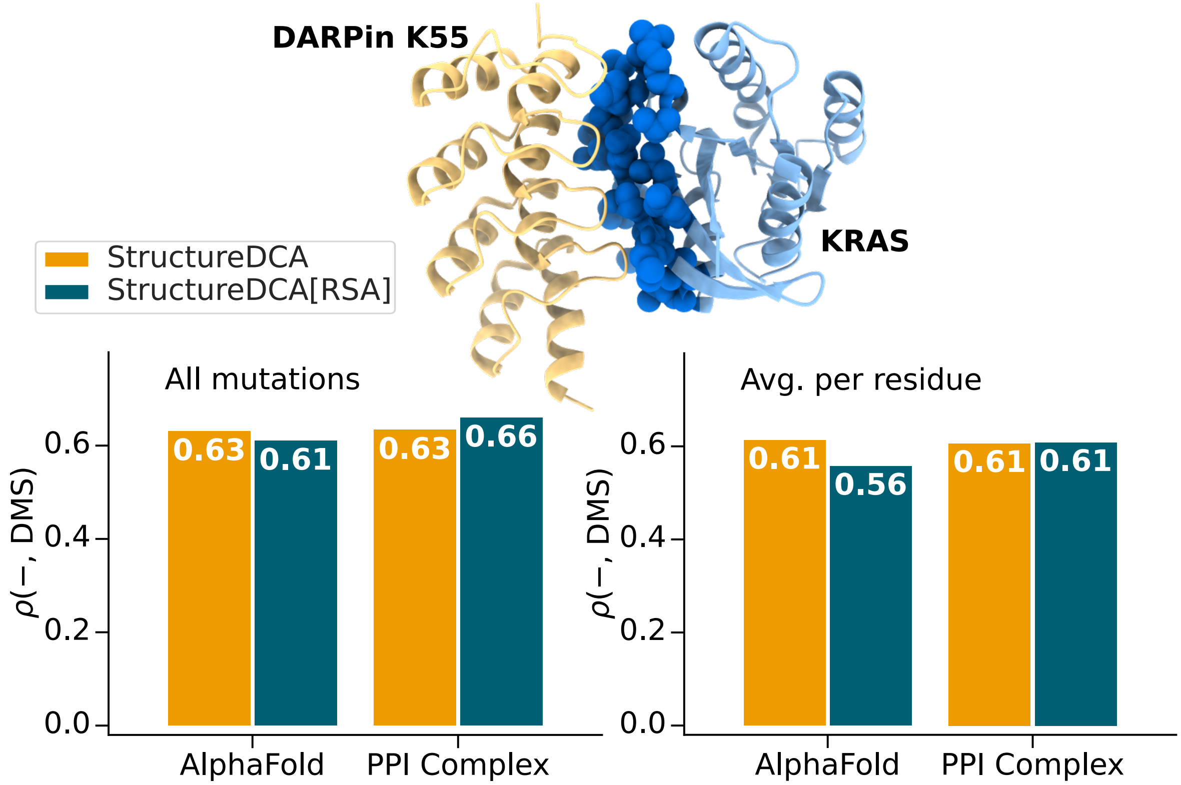

**Figure S13: Interaction between human KRAS and DARPin K55 probed by DMS experiments [33].** Spearman rank correlations of experimental DMS data [33] with StructureDCA and StructureDCA[RSA] predictions, computed using either the AlphaFold monomeric structures or the experimental dimeric structure (PDB code `5mla` [34]). Results are provided for all mutations (left) and for per-residue average values (right). Residues whose RSA is affected by the interaction are highlighted as blue spheres on the 3D structure. The use of the structure of the KRAS-DARPin K55 complex improves performance of StructureDCA[RSA] by about $\rho = 0.05$.
